# Optimizing auditory brainstem response acquisition using interleaved frequencies

**DOI:** 10.1101/781831

**Authors:** Brad N. Buran, Sean Elkins, J. Beth Kempton, Edward V. Porsov, John V. Brigande, Stephen V. David

## Abstract

Auditory brainstem responses (ABRs) require averaging responses to hundreds or thousands of repetitions of a stimulus (e.g., tone pip) to obtain a measurable evoked response at the scalp. Fast repetition rates lead to changes in ABR amplitude and latency due to adaptation. To minimize the effect of adaptation, stimulus rates are sometimes as low as 10 to 13.3 stimuli per second, requiring long acquisition times. The trade-off between reducing acquisition time and minimizing the effect of adaptation on ABR responses is an especially important consideration for studies of cochlear synaptopathy, which use the amplitude of short latency responses (wave 1) to assess auditory nerve survival. It has been proposed that adaptation during ABR acquisition can be reduced by interleaving tones at different frequencies, rather than testing each frequency serially. With careful ordering of frequencies and levels in the stimulus train, adaptation in the auditory nerve can be minimized, thereby permitting an increase in the rate at which tone bursts are presented. However, widespread adoption of this stimulus design has been hindered by lack of available software. Here, we develop and validate an interleaved stimulus design to optimize the rate of ABR measurement while minimizing adaptation. We implement this method in an open-source data acquisition software tool that permits either serial or interleaved ABR measurements. The open-source software library, psiexperiment, is compatible with widely-used ABR hardware. Consistent with previous studies, careful design of an interleaved stimulus train can reduce ABR acquisition time by more than half, with minimal effect on ABR thresholds and wave 1 latency, while improving measures of wave 1 amplitude.

## 1 Introduction

Auditory brainstem responses (ABRs) are an essential tool for assessing peripheral auditory function in animals and humans. In addition to diagnosing peripheral injuries, the ABR is used in newborn hearing screens and intraoperative monitoring of auditory function (Hood, 1998; Burkard et al., 2007). The ABR is measured using electrodes at the scalp and represents the far-field potential of the auditory nerve and several brainstem nuclei (Melcher and Kiang, 1996). Average responses to hundreds or thousands of presentations of a tone pip or click are needed to obtain a measurable evoked response at the scalp. Because neural activity adapts during repeated sensory stimulation, sensitive ABR measurements may require presenting stimuli at rates as low as 10 per second. Faster rates lead to changes in ABR amplitude and latency due to adaptation (Mouney et al., 1976; Paludetti et al., 1983).

The trade-off between reducing ABR acquisition time and minimizing the effect of adaptation is an important consideration for some experiments. For example, studies of moderate noise exposure have found that ABR wave 1 amplitude is a sensitive measure of auditory nerve survival in animals (Furman et al., 2013; Lin et al., 2011; Kujawa and Liberman, 2009, 2006), and this measure has been used as an indirect assessment of hidden hearing loss in humans (Bramhall et al., 2017). Wave amplitude and latency are also of interest clinically in intraoperative monitoring as well as diagnosis of retrocochlear disorders (Hood, 1998; Burkard et al., 2007). Although it is desirable to maximize measures of wave 1 amplitude, many studies of auditory peripheral function in animals use presentation rates that drive adaptation of auditory nerve fibers in order to limit acquisition time (Spoor and Eggermont, 1971; Harris and Dallos, 1979; Burkard and Voigt, 1990). In human studies, slow presentation rates of 10 to 13 tones/s are used when adaptation needs to be minimized (e.g., Bramhall et al., 2017; Stamper and Johnson, 2015). This requirement limits the number of frequencies and levels that can be tested in a reasonable amount of time.

A number of studies have explored various time-saving strategies for ABR measurement. One approach is to randomize the stimulus timing, which allows for faster presentation rates since temporally overlapping responses of adjacent stimuli average out with appropriate randomization of the interstimulus interval (e.g., Eysholdt and Schreiner, 1982; Polonenko and Maddox, 2019; Millan et al., 2006; Valderrama et al., 2012; Burkard et al., 1990; Burkard, 1991). However, studies using this approach have shown evidence of adaptation, manifested as a decrease in ABR amplitude or increase in ABR latency. Recognizing this issue, Mitchell et al. (1999; 1996) designed a novel approach to minimize adaptation by interleaving different stimuli, which took advantage of the tonotopic tuning of auditory nerve fibers. Auditory nerve fibers have sharp frequency tuning (Fig. 1), particularly at low stimulus levels, and do not respond robustly to frequencies above their characteristic frequency (Kiang, 1965) due to the asymmetric spread of basilar membrane excitation towards the apex (i.e., low-frequency region) of the cochlea (Robles and Ruggero, 2001). Thus, careful ordering of the frequencies and levels in the stimulus train can minimize adaptation while increasing the presentation rate. A test of the interleaved stimulus design in normal-hearing humans demonstrated a small, but significant, increase in wave 5 latencies and slight decrease in wave 5 amplitudes (other waves were not analyzed, Henry et al., 2000; Fausti et al., 1994). Attempts to leverage this strategy in monitoring of patients with cisplatin-induced ototoxicity had limited success, potentially because modifications to the interleaved protocol were not first validated in normal-hearing subjects (Dille et al., 2013; Mitchell et al., 2004).

**Fig. 1.**
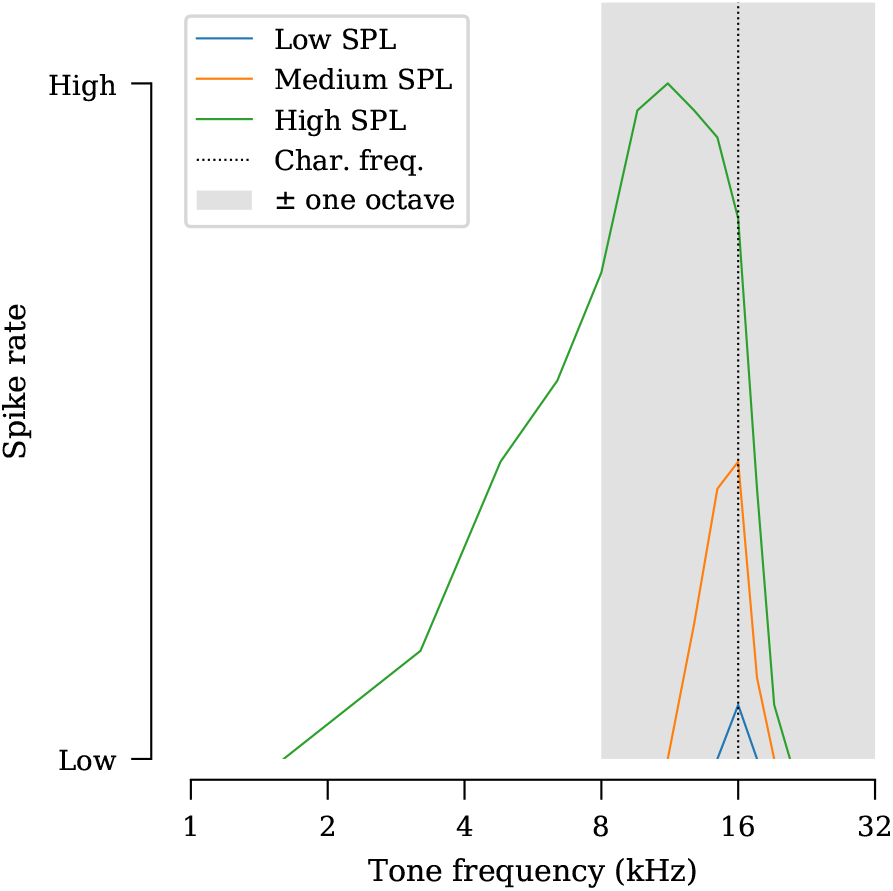
Frequency tuning curve for a single auditory nerve fiber with a characteristic frequency of 16 kHz. To illustrate the asymmetric spread of excitation towards the low-frequency region of the cochlea, tuning curves for low, medium and high-intensity tones are shown. For this fiber, playing a 32 kHz tone immediately prior should minimally affect the response of the fiber to a 16 kHz tone. Shaded area indicates the frequency range from one octave below to one octave above the fiber’s characteristic frequency. Based on data from Kiang (1965).

Given renewed interest in the ABR wave 1 amplitude as a measure of cochlear dysfunction, this study provides additional evidence that ABR studies in animals can benefit from an interleaved stimulus design. We test three parametric manipulations to the interleaved stimulus design that highlight the possible options (e.g., maximizing wave amplitude vs. maximizing acquisition speed). We first confirm that an interleaved stimulus design results in less adaptation as compared to the conventional stimulus design when using the same presentation rate. We then demonstrate that interleaving five frequencies at a rate of 50 tones/s results in ABR amplitudes that are equivalent to those acquired using a conventional approach at a slower rate that does not drive adaptation (10 tones/s). Finally, we demonstrate that optimizing the ordering of frequencies and levels in the interleaved stimulus train yields additional increases in wave amplitude. We tested this approach over a stimulus frequency range of 2 to 32 kHz using mouse and gerbil as our primary model systems. Corroborating data was generated in ferret and rhesus macaque.

One reason why the interleaved approach has not been widely adopted is likely due to hardware and software limitations in most ABR measurement systems. To facilitate use of the interleaved stimulus design, we have written open-source data acquisition software for auditory experiments that implements both the conventional and interleaved stimulus designs described in this paper (Buran and David, 2018). This software runs on the same National Instruments hardware as the widely-used Eaton Peabody Laboratories Cochlear Function Test Suite (Hancock et al., 2015) and is thus readily available to many research groups.

## 2 Materials and Methods

### 2.1 Subjects

All procedures were performed in compliance with the Institutional Animal Care and Use Committee of Oregon Health & Science University and the Office of Laboratory Animal Welfare, Office of Extramural Research, National Institutes of Health.

The majority of auditory brainstem response (ABR) data were acquired from Mongolian gerbils (*Meriones unguiculatus*) and mice (*Mus musculus*). Data from ferret (*Mustela putorius*) and rhesus macaque (*Macaca mulatta*) are also shown for a more limited set of experimental conditions. The number of animals used is reported in the results on a per-experiment basis. Gerbils of either sex were used and spanned an age range of 8 to 16 weeks. Mice of either sex were used and spanned an age range of 4 to 20 weeks. For mouse, some data were from mice of the FVB strain and other data were from heterozygous Ush1C^216GA^ mice (Lentz et al., 2010). The ferret was a three year old spayed and descented male. Rhesus macaques were 5 month old females.

Animals were anesthetized (gerbil: 100 mg/kg ketamine and 10 mg/kg xylazine; mouse: 65 mg/kg ketamine, 6 mg/kg xylazine and 1 mg/kg acepromazine; macaque: 10 mg/kg ketamine and 15 *μ*g/kg dexmedetomidine; ferret: 5 mg/kg ketamine and 0.05 mg/kg dexmedetomidine). Three electrodes were inserted (gerbil and ferret: vertex and pinna with ground near the base of the tail; mouse: vertex and along the ipsilateral mandible with ground in the forepaw; macaque: midline halfway between the forebrow and the vertex of the skull with reference on the mandible ventral to the ear and ground in the shoulder). ABRs were evoked with tone pips. The voltage difference between pinna and vertex was amplified (gerbil and ferret: 100,000×; mouse and macaque: 10,000×), filtered (gerbil and ferret: 0.1 to 10 kHz, mouse and macaque: 0.3 to 3 kHz) and raw traces were digitized for subsequent analysis. For mouse, gerbil and ferret, an Astro-Med Grass P511 amplifier was used. For rhesus macaque, a Signal Recovery Model 5113 amplifier was used. Body temperature was maintained between 36 and 37°C using a homeothermic blanket (ferret, gerbil, mouse) or chemical heat packs (macaque).

Although longer-lasting anesthetics (e.g., isoflurane) are available, these alternatives may have an adverse impact on auditory function (Cederholm et al., 2012; Ruebhausen et al., 2012; Smith and Mills, 1989).

### 2.2 Stimulus Design

Acoustic stimuli were digitally generated (PXI data acquisition system with 24-bit analog-to-digital and digital-to-analog converter PXI-4461 card, National Instruments, Austin, TX) and amplified (SA1, Tucker-Davis Technologies, Alachua, FL). Due to the small size of the mouse ear, a compact, closed-field sound system consisting of two half-inch dome tweeters and an electret microphone (Knowles FG-23329-P07) coupled to a probe tube, was used to deliver acoustic stimuli to the ear. This acoustic system was designed at Eaton-Peabody Laboratories (Hancock et al., 2015) and is colloquially referred to as a starship due to its appearance (Fig. 2). Since the starship cavity alters the frequency response of the electret microphone, the electret microphone was calibrated between 0.1 and 100 kHz using a 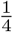 inch microphone that has a flat frequency response (377B10 coupled with 426B03 preamplifier and 480M122 signal conditioner, PCB Piezotronics, Depew, NY). Once the probe tube of the starship was positioned over the tragus of the ear canal, the speakers were calibrated using the probe tube microphone immediately prior to the start of the experiment.

**Fig. 2.**
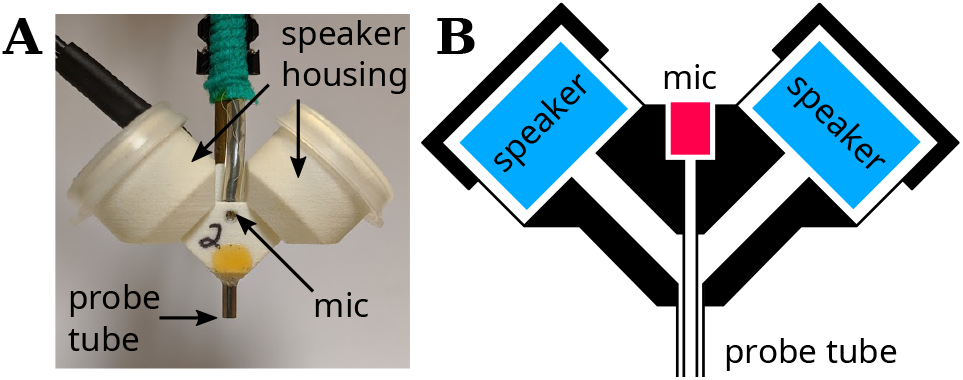
Acoustic system designed by Eaton-Peabody Laboratories for assessing auditory function in animals. **A)** Photograph of the enclosure. The tip of the probe tube is placed immediately next to the ear canal to deliver sound. **B)** Schematic showing a cross-section of the enclosure, which contains two speakers (blue) and a microphone (pink). Only one speaker is required for ABR experiments.

ABRs were generated using 5 ms tone pips with an 0.5 ms cosine-squared envelope. Levels were incremented in 5 dB steps from 10 to 80 dB SPL. The order of tone pip presentation depended on the stimulus design (Fig. 3):

- **conventional**: Tone pips were repeated at a fixed frequency and level until the desired number of artifact-free trials was acquired (Fig. 3A).
- **interleaved**: A train of tone pips containing a single presentation of each level and frequency was constructed. This train was then presented repeatedly until the desired number of artifact-free trials was acquired (Fig. 3B-D).

**Fig. 3.**
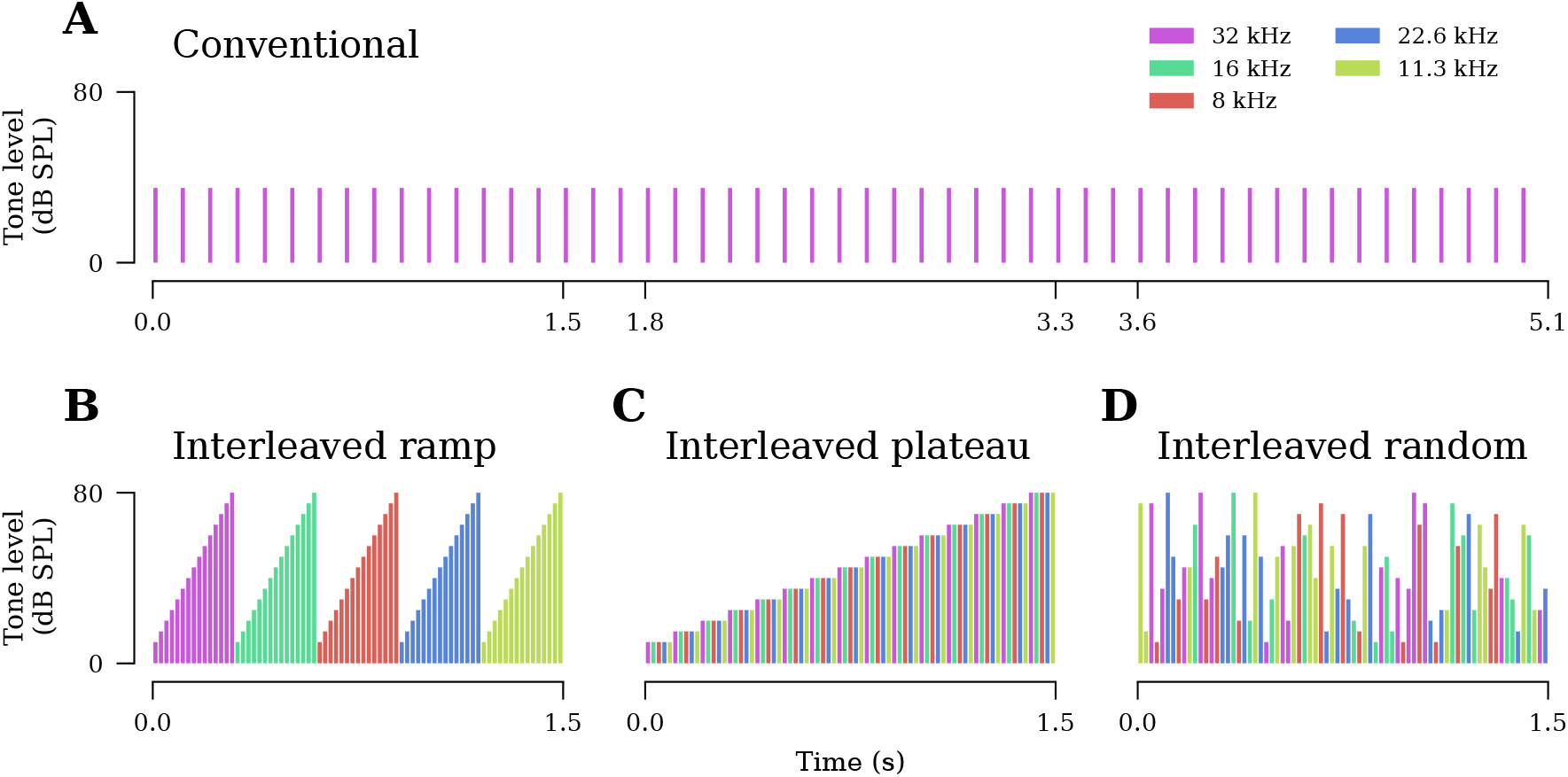
Schematic of the conventional and interleaved ABR stimulus designs tested. **A)** The conventional approach presents tone pips at a single frequency and level until the desired number of averages are acquired. **B-D)** In contrast, the interleaved approach presents a stimulus train containing a single tone pip of each tested frequency and level. This train is then presented repeatedly until the desired number of averages are acquired. For details regarding the ordering of frequencies and levels in each interleaved protocol, see text. Note the horizontal time axis is plotted at the same scale for all four sequences, highlighting differences in tone presentation rate, 10 tones/s for conventional and 50 tones/s for interleaved. These rates were used for data shown in Fig. 6.

For the interleaved stimulus designs, the presentation rate was defined as the rate at which individual tones appear in the train (i.e., a 5 frequency, 15 level train with a presentation rate of 50 tones/s would result in a 1.5 second long train). We tested three different rules for constructing the interleaved stimulus train:

- **ramp**: Levels were swept from low to high before advancing to the next frequency (Fig. 3B).
- **plateau**: All frequencies were presented at a fixed level before advancing to the next higher level (Fig. 3C).
- **random**: The set of levels and frequencies was shuffled randomly on each presentation of the train (Fig. 3D).

In the conventional stimulus design, the polarity of tone pips was alternated on each presentation of the tone pip to remove frequency-following responses. In the interleaved stimulus designs, the polarity of all tone pips was alternated between each presentation of a train.

All interleaved experiments in mouse and gerbil tested a sequence of five frequencies. For the interleaved ramp and interleaved plateau designs, frequencies were arranged in decreasing order while maintaining a minimum spacing of one octave between adjacent frequencies. The exact order was 8, 4, 2, 5.7 and 2.8 kHz for gerbil and 32, 16, 8, 22.6 and 11.3 kHz for mouse. For ferret, we acquired 2 to 45.2 kHz in half-octave steps using the interleaved random stimulus. In macaque, 0.5, 2, 4, 8, 16, 22.6 and 32 kHz were tested using the interleaved ramp stimulus design. Since auditory nerve fibers are preferentially tuned to frequencies within half an octave of the characteristic frequency of the fiber (Kiang, 1965, Fig. 1), tones falling outside of this range should not drive much adaptation of the fiber (Harris and Dallos, 1979).

When comparing two or more stimulus designs (e.g., conventional at 10/s vs. interleaved ramp at 50/s), all permutations were tested within a single animal during a single session (i.e., no re-positioning of the electrodes and/or acoustic system). To avoid biases introduced by variations in anesthesia depth, the ordering of the stimulus designs and presentation rates were randomized for each ear. Anesthetic booster dosing was only necessary when acquiring data from gerbil and care was taken to perform the injection without altering the position of the animal’s head. For mouse, data were acquired without using an anesthetic booster.

### 2.3 Artifact rejection

For interleaved studies, only the segment of the train containing the artifact was rejected, rather than rejecting the entire stimulus train. This means we acquired a variable number of artifact-free averages for each frequency and level tested, but every frequency and level had at least 512 averages. When generating waveforms for analysis, only the first 512 averages were included to ensure the number of averages was identical across all experiments.

### 2.4 Analysis

ABR waveforms were extracted (−1 to 10 ms re tone pip onset) and averaged. To match the filter settings for mice, gerbil waveforms were digitally filtered (0.3 to 3 kHz) prior to averaging. Thresholds were identified via visual inspection of stacked waveforms by two trained observers, each blind to the stimulus design. Results from the two observers were compared and discrepancies of greater than 10 dB reconciled. Wave amplitude and latency were identified using a computer-assisted peak-picking program (Buran, 2015). Wave amplitude was defined as the difference between the peak and the following trough.

#### Mixed linear models

Differences in ABR threshold, wave amplitude and wave latency were assessed using a general mixed linear model. In this model, *y*_*i*_ represents the measured value (ABR threshold, wave amplitude or wave latency). For wave amplitude and latency, the measured value at 80 dB SPL was used. Intercept (*β*_*i*_) allowed for a constant offset. Stimulus frequency (*β*_*f*_), stimulus design (*β*_*c*_) and repetition rate (*β*_*r*_) were fixed effects. All two-way (*β*_*fc*_, *β*_*rf*_ and *β*_*rc*_) and three-way (*β*_*rfc*_) interactions between the fixed effects were included. Frequency, *f*, and stimulus design, *c*, were treated as categorical parameters and rate, *r*, as a continuous parameter. Dummy (i.e., treatment) coding was used for all categorical parameters. Since both mouse and gerbil were tested at 8 kHz, data from each species were coded separately at this frequency to avoid introducing an additional effect for species. Ear was treated as a random effect and coded as *U*_*e*_ where *e* represents index of the ear.

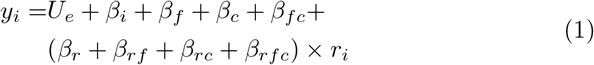

For comparing sequential at 10 tones/s versus interleaved ramp at 50 tones/s, the rate parameters were dropped, simplifying the model to:

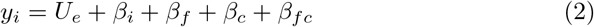

#### Bayesian regression

All models were fit using Bayesian regression to maximize the Normal likelihood of free parameters using pyMC3 (Salvatier et al., 2016). In contrast to conventional model fitting, Bayesian analysis allows for simple calculations of credible intervals on derived parameters (e.g., parameters that are mathematical functions of fitted coefficients), offers simple construction of realistic hierarchical models (Gelman et al., 2013), avoids a number of problems with conventional p-values derived from null hypothesis significance testing (Szucs and Ioannidis, 2017), determines the probability that model coefficients take on a particular value or range of values (McMillan and Cannon, 2019) and allows for accepting the null value when certainty in the estimate is high (Kruschke, 2013).

Diffuse priors were used to ensure minimal influence on the parameter estimates. The intercept, *β*_*i*_, had a Normal prior with a mean and standard deviation set to the mean and standard deviation, respectively, of the pooled data (Kruschke, 2013). All other parameters had a Normal prior with mean of 0 and standard deviation set to the standard deviation of the pooled data. Each model was fit four times for 2000 samples following a 1000 sample burn-in period using the No U-Turn Sampler (Hoffman and Gelman, 2014). Posterior samples were combined across all fits (i.e., chains) for inference. Gelman-Rubin statistics were computed to ensure that the four fits, each of which started with a random estimate for each parameter, converged to the same final estimate 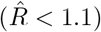.

#### On p-values

Unlike conventional (i.e., frequentist) approaches, Bayesian analysis does not offer p-values. Instead, Bayesian analysis quantifies the probability that the true value for a parameter falls between two points. These distributions can be used to calculate the probability that there is a true difference between groups, which is typically the information people incorrectly attempt to glean from p-values (Nuzzo, 2014). In our analyses, we report the mean and 95% credible interval (CI) for the difference between groups (e.g., interleaved ramp vs. conventional). The CI should be interpreted as the interval in which we are 95% certain contains the true value. Therefore, if the 95% CI does not bracket 0, we can assume the value is significantly different from 0. To further aid in interpretation of our results, we calculate the probability that the difference in the parameter between two groups does not exceed a certain range (e.g., ±2.5 dB) by integrating over the portion of the posterior distribution that falls within that range (see Table 1 caption for an example of interpreting this posterior probability value).

**Table 1.** Change in ABR thresholds in dB relative to 40/s conventional stimulus design. Negative values indicate the threshold was lower than for 40/s conventional. Significance is assessed by the posterior probability that the difference in threshold from 40/s conventional is less than 2.5 or 5 dB; e.g., a probability of 0.97 for 2.5 dB indicates a 97% chance that the difference in ABR threshold is between −2.5 and 2.5dB and a 3% chance that the difference in ABR threshold is less than −2.5 dB or greater than 2.5 dB.

| Stim. design | Species | Freq. | dB re. 40/s conv. |  | Probability |  |
| --- | --- | --- | --- | --- | --- | --- |
| | | | Mean | 95% CI | < $\pm 2.5$ dB | < $\pm 5$ dB |
| 40/s int. ramp | gerbil | 2.8 kHz | -0.8 | ( -2.9, 1.3) | 0.94 | 1.00 |
|  |  | 5.7 kHz | -0.7 | ( -3.0, 1.5) | 0.94 | 1.00 |
|  | mouse | 11.3 kHz | -1.3 | ( -3.9, 1.3) | 0.81 | 1.00 |
|  |  | 22.6 kHz | -2.2 | ( -4.9, 0.5) | 0.60 | 0.98 |
| 80/s int. ramp | gerbil | 2.8 kHz | 0.2 | ( -2.0, 2.6) | 0.97 | 1.00 |
|  |  | 5.7 kHz | -0.3 | ( -2.7, 2.1) | 0.95 | 1.00 |
|  | mouse | 11.3 kHz | -0.7 | ( -3.6, 2.0) | 0.88 | 1.00 |
|  |  | 22.6 kHz | -1.9 | ( -4.7, 0.9) | 0.65 | 0.98 |
| 80/s conv. | gerbil | 2.8 kHz | 1.1 | ( -0.5, 2.8) | 0.95 | 1.00 |
|  |  | 5.7 kHz | -0.1 | ( -2.2, 2.2) | 0.97 | 1.00 |
|  | mouse | 11.3 kHz | 2.5 | ( -0.1, 4.9) | 0.49 | 0.97 |
|  |  | 22.6 kHz | 1.2 | ( -1.3, 3.6) | 0.83 | 1.00 |

## 3 Results

We first test whether ABR data acquired using an interleaved stimulus design show less adaptation than data acquired using a conventional stimulus design at the same presentation rate. Next, we assess whether interleaving five frequencies at a rate of 50 tones/s can produce results equivalent to a conventional approach at a rate of 10 tones/s that does not drive adaptation. Finally, we test whether we can reduce adaptation even further by modifying the order of the stimuli within the interleaved stimulus design.

### 3.1 Conventional vs. interleaved ramp at matched rates

We first assessed whether the interleaved ramp stimulus design offers an advantage over the conventional design for stimuli presented at the same rate. We compared two rates, 40/s, which is common in the animal literature, and 80/s, to assess whether doubling the presentation rate yielded additional benefits. Measurements were compared for both mouse and gerbil. A set of five frequencies in each species was assessed when using the interleaved ramp design, but time constraints from anesthesia duration limited the number of frequencies measured in the conventional protocol (2.8 and 5.7 kHz for gerbil, 11.3 and 22.6 kHz for mouse). Thus, data are shown only for the subset of frequencies common to both stimulus designs.

Both the interleaved ramp and conventional stimulus designs yielded clean ABR waveforms, with waves 1 through 5 easily identifiable (Fig. 4A). At presentation rates of 40/s and 80/s, the interleaved ramp design had ABR thresholds that were at least as low as thresholds acquired using a 40/s conventional design (Fig. 4B, Table 1). Regardless of presentation rate or species, ABR thresholds in the interleaved stimulus designs were within ±5 dB of the conventional stimulus design (Table 1). Wave 1 amplitudes, defined as the difference between the first peak and following trough (Fig. 4A), were larger in both the 40 and 80/s interleaved ramp stimulus design, as compared to 40/s conventional, for all stimulus levels tested (Fig. 4C). In particular, wave 1 amplitudes for 40/s interleaved ramp were 22-26% larger than 40/s conventional and 80/s interleaved ramp were 8-21% larger than 40/s conventional (Fig. 4D, Table 2). In contrast, wave 1 amplitudes in 80/s conventional were 13-23% smaller than 40/s conventional. Waves 2 through 5 had amplitudes in the 40 and 80/s interleaved design that were at least as large as 40/s conventional (Fig. 5A,C,E,G). Although there were some differences in wave latencies, they were less than ±0.2 ms (Figs. 4E,F and 5B,D,F,H, Table 3).

**Fig. 4.**
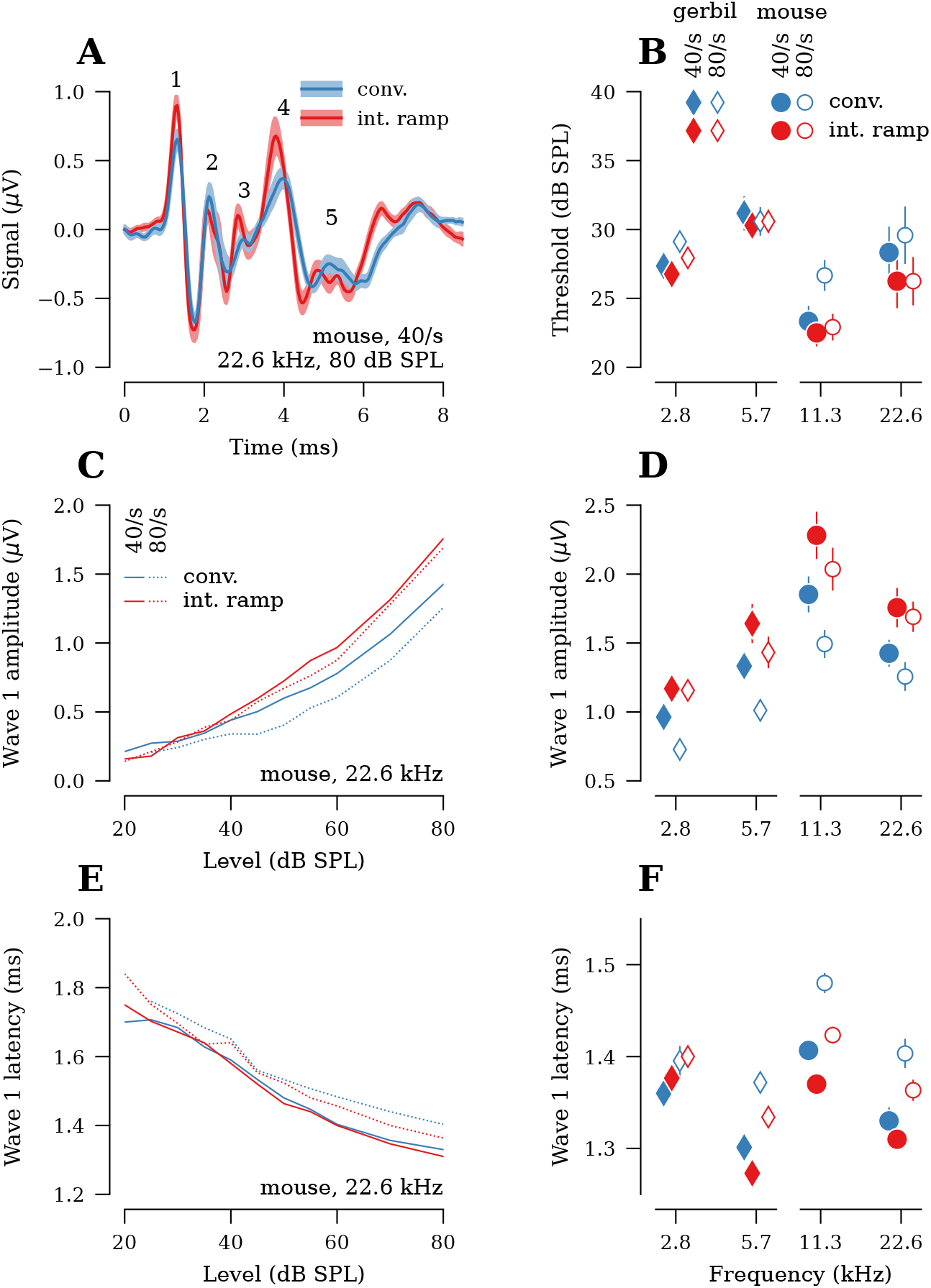
The interleaved ramp stimulus design yields greater wave 1 amplitudes than the conventional stimulus design, regardless of presentation rate (*n*=17 ears from gerbil, 8 ears from mouse). **A)** Comparison of ABR waveforms (ensemble average across all ears) acquired using the interleaved and conventional stimulus designs. Data shown are in response to 40/s 22.6 kHz, 80 dB SPL tone pips in mouse. Numbers indicate wave. **B)** Average ABR thresholds for each frequency, presentation rate and stimulus design. **C)** Average wave 1 amplitudes vs. stimulus level for 40/s 22.6 kHz tones in mouse. **D)** Average wave 1 amplitudes at 80 dB SPL. **E)** Average wave 1 latencies vs. stimulus level for 40/s 22.6 kHz tones in mouse. **F)** Average wave 1 latencies at 80 dB SPL. Shaded area and error bars in all panels indicate ±SEM.

**Table 2.** Percent change in wave 1 amplitudes relative to 40/s conventional stimulus design. Negative values indicate the amplitude measurement was lower than for 40/s conventional. Significance is assessed by the posterior probability that the change in wave 1 amplitude from 40/s conventional is less than ±5 or ±10%.

| Stim. design | Species | Freq. | % re. 40/s conv. |  | Probability |  |
| --- | --- | --- | --- | --- | --- | --- |
| | | | Mean | 95% CI | < $\pm 5\%$ | < $\pm 10\%$ |
| 40/s int. ramp | gerbil | 2.8 kHz | 22 | ( 6, 40) | 0.01 | 0.07 |
|  |  | 5.7 kHz | 23 | ( 10, 37) | 0.00 | 0.02 |
|  | mouse | 11.3 kHz | 26 | ( 12, 39) | 0.00 | 0.01 |
|  |  | 22.6 kHz | 26 | ( 10, 44) | 0.00 | 0.02 |
| 80/s int. ramp | gerbil | 2.8 kHz | 19 | ( 2, 36) | 0.04 | 0.15 |
|  |  | 5.7 kHz | 8 | ( -4, 21) | 0.31 | 0.63 |
|  | mouse | 11.3 kHz | 12 | ( 1, 25) | 0.12 | 0.37 |
|  |  | 22.6 kHz | 21 | ( 3, 38) | 0.02 | 0.09 |
| 80/s conv. | gerbil | 2.8 kHz | -23 | (-36, -10) | 0.00 | 0.02 |
|  |  | 5.7 kHz | -23 | (-34, -12) | 0.00 | 0.01 |
|  | mouse | 11.3 kHz | -20 | (-31, -10) | 0.00 | 0.03 |
|  |  | 22.6 kHz | -13 | (-27, 2) | 0.12 | 0.33 |

**Fig. 5.**
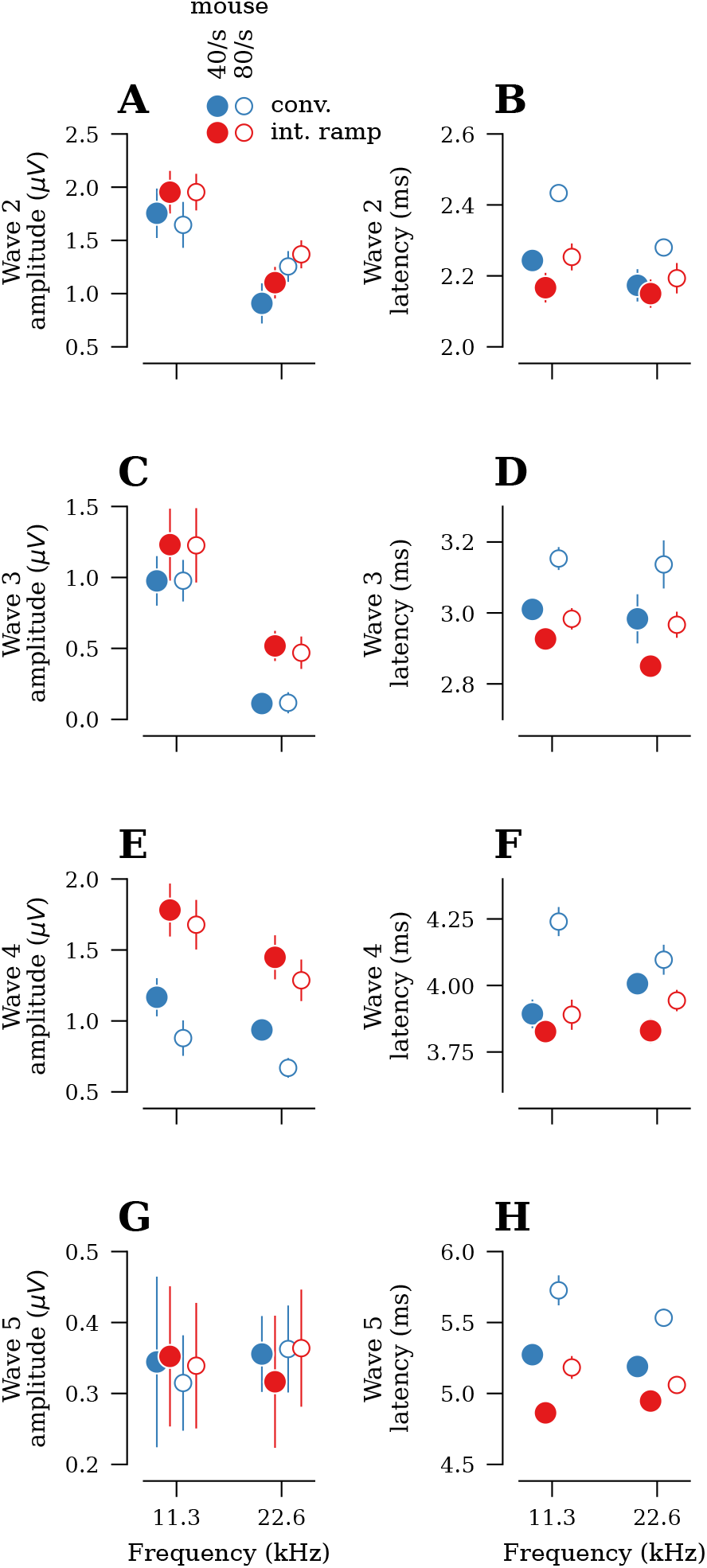
Comparison of ABR wave 2-5 amplitudes and latencies, acquired using conventional and interleaved ramp stimulus designs at all frequencies and presentation rates tested (see Fig. 4 for wave 1 data; *n* = 8 ears in mouse). **A,C,E,G)** Average wave amplitudes at 80 dB SPL. **B,D,F,H)** Average wave latencies at 80 dB SPL. Error bars in all panels indicate ±SEM.

**Fig. 6.**
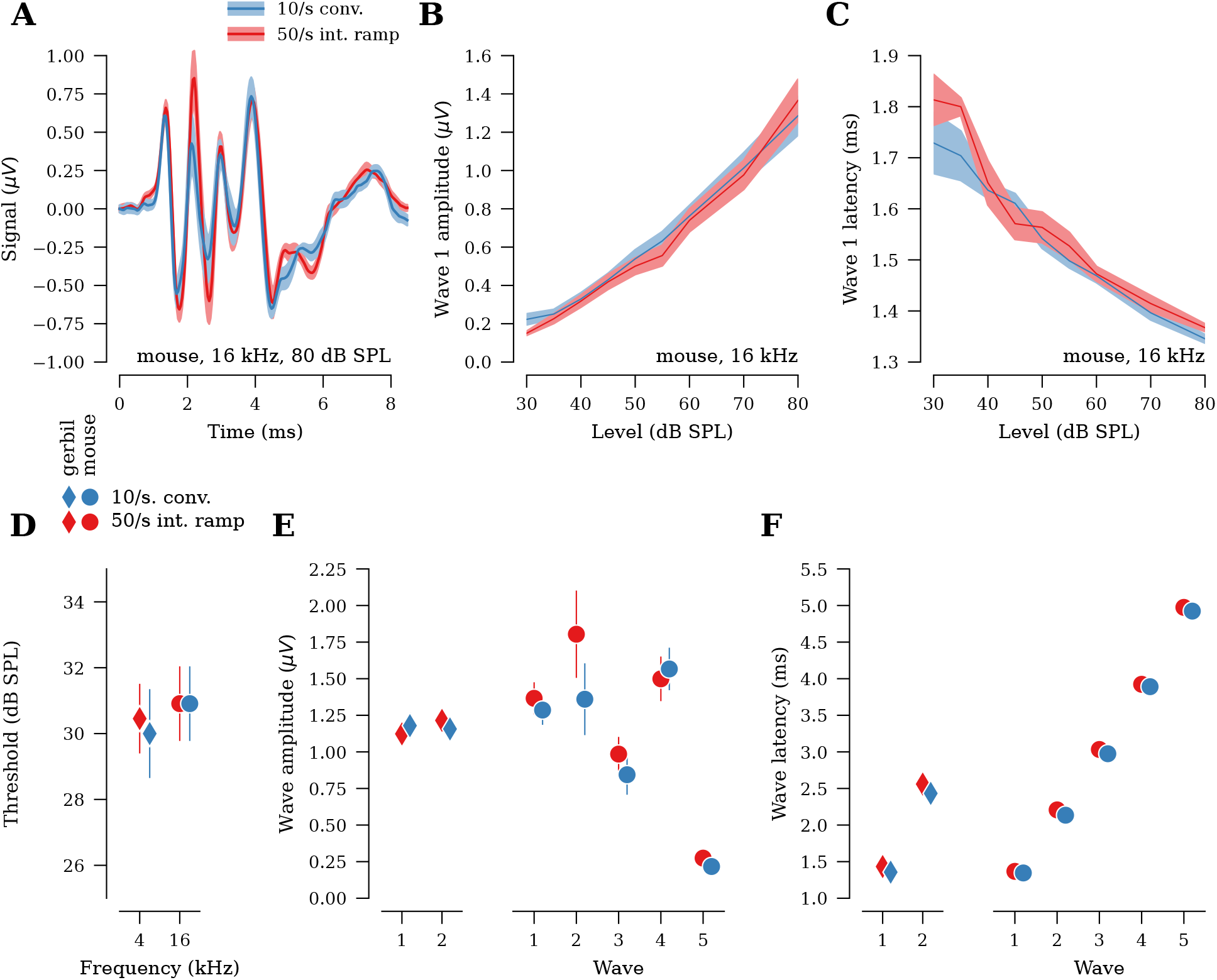
ABR metrics acquired using a five-frequency 50/s interleaved ramp protocol are comparable to a 10/s conventional protocol for both low (4 kHz) and high (16 kHz) frequencies (*n* = 11 ears from gerbil, *n* = 11 ears from mouse). **A)** ABR waveforms (ensemble average across all ears) in response to 80 dB SPL tone pips at 16 kHz in mouse. **B)** Average wave 1 amplitudes vs. stimulus level for 16 kHz tones in mouse. **C)** Average wave 1 latencies vs. stimulus level for 16 kHz tones in mouse. **D)** Average ABR thresholds. **E)** Average wave amplitudes at 80 dB SPL. **F)** Average wave latencies at 80 dB SPL. Shaded area and error bars in all panels indicate ±SEM. Error bars in F are too small to be visible.

**Table 3.** Change in wave 1 latencies in ms relative to 40/s conventional stimulus design. Negative values indicate the latency was shorter than for 40/s conventional. Significance is assessed by the posterior probability that the difference in wave 1 latency from 40/s conventional is less than ±0.1 or ±0.2 ms.

| Stim. design | Species | Freq. | ms re. 40/s conv. |  | Probability |  |
| --- | --- | --- | --- | --- | --- | --- |
| | | | Mean | 95% CI | < $\pm 0.1$ ms | < $\pm 0.2$ ms |
| 40/s int. ramp | gerbil | 2.8 kHz | 0.01 | (-0.01, 0.04) | 1.00 | 1.00 |
|  |  | 5.7 kHz | -0.03 | (-0.05, -0.01) | 1.00 | 1.00 |
|  | mouse | 11.3 kHz | -0.04 | (-0.07, -0.01) | 1.00 | 1.00 |
|  |  | 22.6 kHz | -0.02 | (-0.05, 0.01) | 1.00 | 1.00 |
| 80/s int. ramp | gerbil | 2.8 kHz | 0.04 | ( 0.02, 0.07) | 1.00 | 1.00 |
|  |  | 5.7 kHz | 0.03 | ( 0.00, 0.05) | 1.00 | 1.00 |
|  | mouse | 11.3 kHz | 0.01 | (-0.01, 0.04) | 1.00 | 1.00 |
|  |  | 22.6 kHz | 0.03 | ( 0.00, 0.06) | 1.00 | 1.00 |
| 80/s conv. | gerbil | 2.8 kHz | 0.04 | ( 0.02, 0.06) | 1.00 | 1.00 |
|  |  | 5.7 kHz | 0.07 | ( 0.04, 0.09) | 1.00 | 1.00 |
|  | mouse | 11.3 kHz | 0.07 | ( 0.04, 0.10) | 0.99 | 1.00 |
|  |  | 22.6 kHz | 0.07 | ( 0.04, 0.09) | 0.99 | 1.00 |

Taken together, these results demonstrate that an interleaved ramp design that doubles presentation rate to 80 tones/s yields ABR results that are equivalent to, and often of greater amplitude than, results acquired using a conventional design at 40 tones/s with minimal effect on ABR threshold and wave latency.

### 3.2 Conventional at 10/s vs. interleaved ramp at 50/s

Slower presentation rates of 10 to 13/s are sometimes used in research studies (e.g., Bramhall et al., 2017) that seek to minimize effects of adaptation that occur at rates above 20/s (Fowler and Noffsinger, 1983; Paludetti et al., 1983). Routine measurements at these slow rates are typically not feasible in anesthetized animals since prolonged anesthesia can have adverse effects on the subjects’ metabolism. Here, we assess whether the interleaved ramp design allows rapid acquisition of ABR data equivalent to that acquired using a conventional stimulus design at 10/s. Since we used five frequencies in the interleaved ramp protocol, a rate of 50/s results in an effective rate of 10/s for each frequency.

We measured ABRs at one frequency using the slower-rate conventional stimulus design (4 kHz in gerbil, 16 kHz in mouse) and compared results to those for the same tone frequency using the 50/s interleaved design (Fig. 6A-C). ABR thresholds between the two stimulus designs were within ±2.5 dB (Fig. 6D, Table 4). Wave 1 amplitudes for the 50/s interleaved ramp design were 5% smaller in gerbil and 6% greater in mouse as compared to 10/s conventional (Fig. 6E, Table 5). Despite the apparent decrease in wave 1 amplitudes for gerbil, the 95% CI bracketed 0% (i.e., no change) and there was a 90% probability that the difference in wave 1 amplitudes were less than ±10%. Wave latencies were similar between the designs (Fig. 6F, Table 6).

**Table 4.** Change in ABR thresholds for 50/s interleaved ramp in dB relative to 10/s conventional stimulus design. Negative values indicate the threshold for 50/s ramp was lower than for 10/s conventional. Significance is assessed by the posterior probability that the difference in threshold from 10/s conventional is less than ±2.5 or ±5 dB.

| Species | dB re. 10/s conv. |  | Probability |  |
| --- | --- | --- | --- | --- |
| | Mean | 95% CI | $< \pm 2.5$ dB | $< \pm 5$ dB |
| gerbil | 0.4 | (-1.8, 2.5) | 0.97 | 1.00 |
| mouse | 0.0 | (-2.2, 2.2) | 0.97 | 1.00 |

**Table 5.**
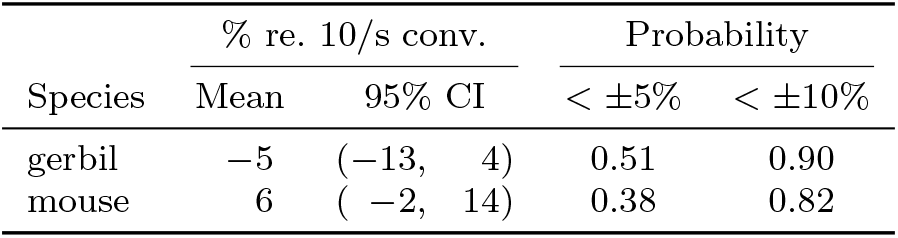
Percent change in wave 1 amplitudes for 50/s interleaved ramp relative to 10/s conventional. Negative values indicate the amplitude for 50/s ramp was lower than for 10/s conventional. Significance is assessed by the posterior probability that the change in wave 1 amplitude from 10/s conventional is less than ±5 or ±10%.

| Species | % re. 10/s conv. |  | Probability |  |
| --- | --- | --- | --- | --- |
| | Mean | 95% CI | $< \pm 5\%$ | $< \pm 10\%$ |
| gerbil | -5 | (-13, 4) | 0.51 | 0.90 |
| mouse | 6 | (-2, 14) | 0.38 | 0.82 |

**Table 6.** Change in wave 1 latencies for 50/s interleaved ramp in ms relative to 10/s conventional. Negative values indicate the latency for 50/s ramp was shorter than for 10/s conventional. Significance is assessed by the posterior probability that the difference in wave 1 latency from 10/s conventional is less than ±0.1 and ±0.2 ms.

| Species | ms re. 10/s conv. |  | Probability |  |
| --- | --- | --- | --- | --- |
| | Mean | 95% CI | $< \pm 0.1$ ms | $< \pm 0.2$ ms |
| gerbil | 0.07 | (0.04, 0.10) | 0.97 | 1.00 |
| mouse | 0.02 | (-0.01, 0.05) | 1.00 | 1.00 |

Taken together, these results demonstrate that using an interleaved ramp design with a presentation rate of 50 tones/s minimizes adaptation of auditory nerve responses and produces results nearly identical to ABR measurements using a 10 tone/s conventional design.

### 3.3 Refining the ordering of interleaved stimuli

In the experiments described above, the interleaved ramp design grouped stimuli by frequency within a single stimulus train (Fig. 3B). This grouping may still drive some adaptation since it rapidly sweeps through the sequence of levels for a single frequency before advancing to the next frequency. To test whether we can obtain further improvements, we assessed two alternative approaches to stimulus ordering: interleaved plateau and interleaved random (Fig. 3C,D). The interleaved plateau design groups stimuli by level, thereby sweeping through all frequencies at a particular level before moving to the next higher level. In this design, tones at each frequency are presented at 20% of the overall rate. For example, a 16 kHz tone in a five-frequency, 80 tones/s interleaved plateau train would appear at a rate of only 16 tones/s. However, at the highest stimulus levels there may be some overlap in excitation patterns along the cochlear partition (Fig. 1; Kiang, 1965; Robles and Ruggero, 2001). The interleaved random design orders tone frequency and level randomly. It imposes no constraints on the grouping of stimuli and may offer a compromise between the interleaved ramp (grouped by frequency) and interleaved plateau (grouped by level) stimulus designs.

To emphasize possible differences between the three stimulus designs, we measured the ABR using a presentation rate of 80 tones/s (Fig. 7A). There was no difference in response thresholds between the three stimulus designs, with the exception that thresholds for interleaved random were 1.9-4.3 dB greater than ramp (Fig. 7B, Table 7). Even for the frequency with the greatest increase in threshold (4.3 dB; 11.3 kHz), the probability that the threshold exceeded ±5 dB was only 30%. ABR wave 1 amplitudes were up to 38% greater in random compared to ramp, although the size of the difference varied quite a bit by frequency (Fig. 7C-D, Table 8). Interleaved plateau generally yielded smaller wave 1 amplitudes than ramp, but the size of the difference varied by frequency. Results for later waves were more variable (Fig. 8A,C,E,G). For latencies, all waves generally occurred slightly earlier in both random and plateau relative to ramp (Fig. 7E-F, Fig. 8B,D,F,H, Table 9).

**Fig. 7.**
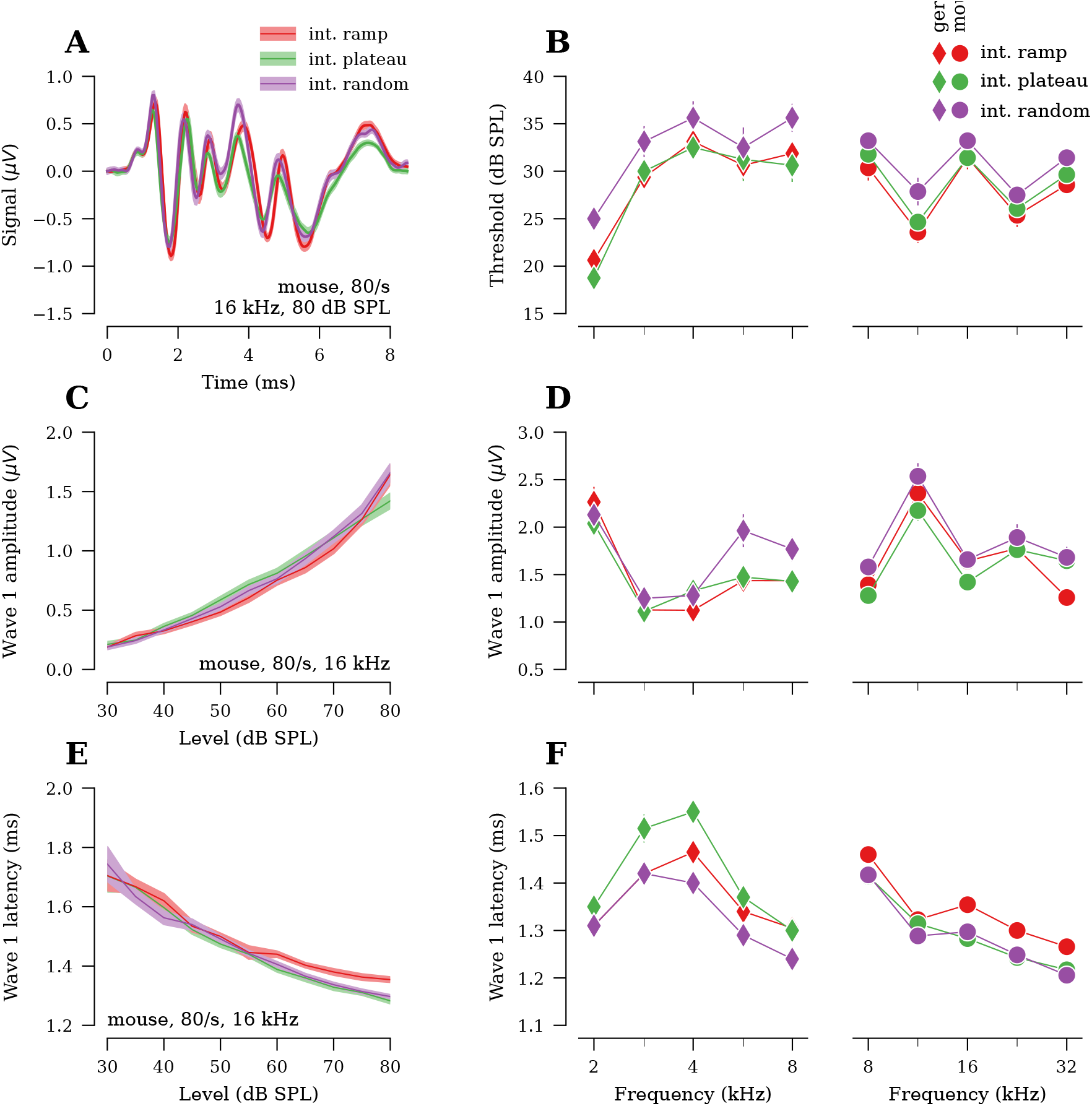
Comparison of interleaved stimulus designs (*n* = 8 ears from gerbil, *n* = 10 ears from mouse). **A)** ABR waveforms (ensemble average across all ears) for different stimulus designs in response to 80/s 16 kHz, 80 dB SPL tone pips in mouse. **B)** Average ABR thresholds. **C)** Average wave 1 amplitudes vs. stimulus level for 80/s 16 kHz tones in mouse. **D)** Average wave 1 amplitudes at 80 dB SPL. **E)** Average wave 1 latencies vs. stimulus level for 80/s 16 kHz tones in mouse. **F)** Average wave 1 latencies at 80 dB SPL. Shaded area and error bars in all panels indicate ±SEM.

**Table 7.** Change in ABR thresholds for 80/s interleaved plateau and interleaved random stimulus designs in dB relative to 80/s interleaved ramp. Negative values indicate the threshold was lower than for 80/s interleaved ramp. Significance is assessed by the posterior probability that the difference in threshold from interleaved ramp is less than ±2.5 or ±5 dB.

| Stim. design | Species | Freq. | dB re. 80/s int. ramp |  | Probability |  |
| --- | --- | --- | --- | --- | --- | --- |
| | | | Mean | 95% CI | $< \pm 2.5$ dB | $< \pm 5$ dB |
| int. plateau | gerbil | 2.0 kHz | -3.4 | (-6.4, -0.0) | 0.30 | 0.83 |
|  |  | 2.8 kHz | 0.7 | (-2.7, 4.2) | 0.82 | 0.99 |
|  |  | 4.0 kHz | -0.5 | (-3.8, 3.1) | 0.83 | 0.99 |
|  |  | 5.7 kHz | 0.7 | (-2.8, 4.0) | 0.82 | 0.99 |
|  |  | 8.0 kHz | -1.1 | (-4.4, 2.1) | 0.78 | 0.99 |
|  | mouse | 8.0 kHz | 1.4 | (-1.1, 4.0) | 0.79 | 1.00 |
|  |  | 11.3 kHz | 1.1 | (-1.6, 3.7) | 0.85 | 1.00 |
|  |  | 16.0 kHz | 0.1 | (-2.4, 2.9) | 0.94 | 1.00 |
|  |  | 22.6 kHz | 0.7 | (-1.7, 3.5) | 0.90 | 1.00 |
|  |  | 32.0 kHz | 1.1 | (-1.6, 3.6) | 0.85 | 1.00 |
| int. random | gerbil | 2.0 kHz | 2.6 | (-0.5, 5.9) | 0.47 | 0.93 |
|  |  | 2.8 kHz | 3.8 | (0.2, 7.1) | 0.22 | 0.75 |
|  |  | 4.0 kHz | 2.6 | (-0.7, 6.1) | 0.47 | 0.92 |
|  |  | 5.7 kHz | 2.0 | (-1.4, 5.3) | 0.61 | 0.96 |
|  |  | 8.0 kHz | 3.8 | (0.5, 7.4) | 0.22 | 0.76 |
|  | mouse | 8.0 kHz | 2.9 | (0.4, 5.6) | 0.37 | 0.94 |
|  |  | 11.3 kHz | 4.3 | (1.7, 6.8) | 0.08 | 0.70 |
|  |  | 16.0 kHz | 1.9 | (-0.6, 4.5) | 0.68 | 0.99 |
|  |  | 22.6 kHz | 2.2 | (-0.4, 4.8) | 0.59 | 0.98 |
|  |  | 32.0 kHz | 2.9 | (0.5, 5.7) | 0.38 | 0.94 |

**Table 8.**
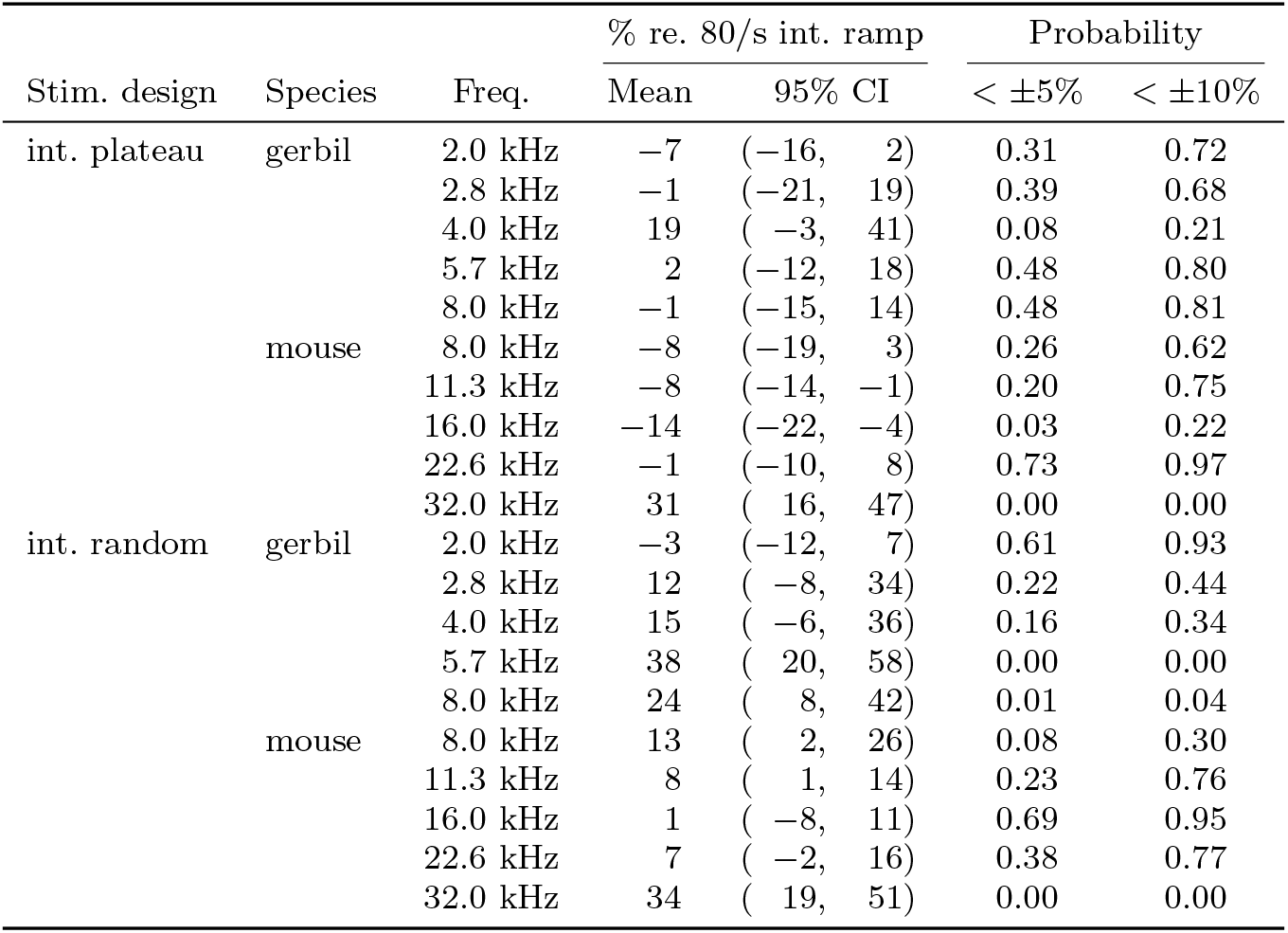
Percent change in wave 1 amplitudes for 80/s interleaved plateau and interleaved random stimulus designs relative to 80/s interleaved ramp. Negative values indicate the amplitude measurement was lower than for 80/s interleaved ramp. Significance is assessed by the posterior probability that change in wave 1 amplitude from 80/s interleaved ramp is less than ±5 or ±10%.

**Fig. 8.**
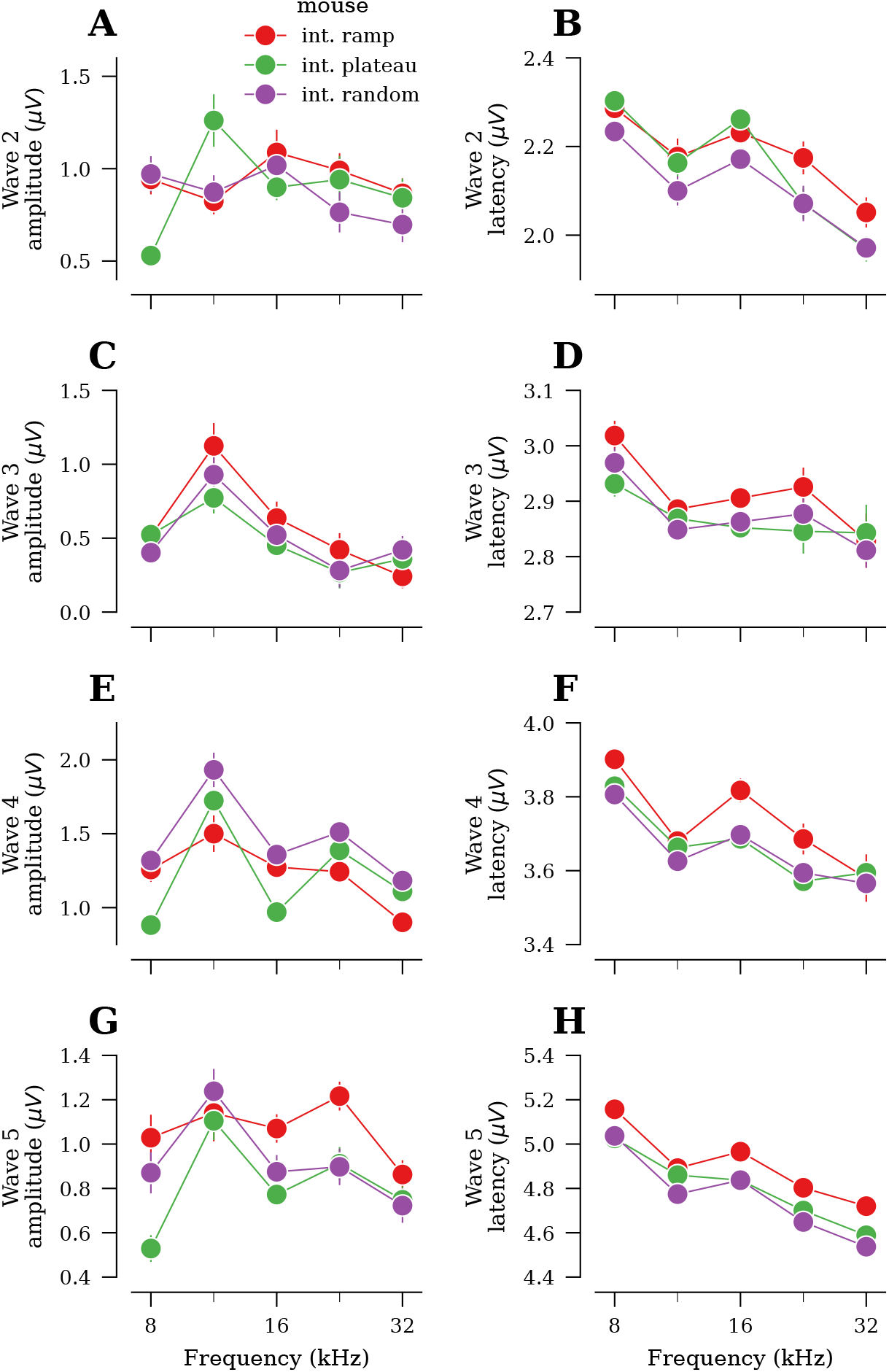
Comparison of ABR wave 2-5 amplitudes and latencies acquired using different interleaved stimulus designs for all frequencies tested (see Fig. 7 for wave 1 data; *n* = 8 ears in mouse). **A,C,E,G)** Average wave amplitudes at 80 dB SPL. **B,D,F,H)** Average wave latencies at 80 dB SPL. Error bars in all panels indicate ±SEM.

**Table 9.**
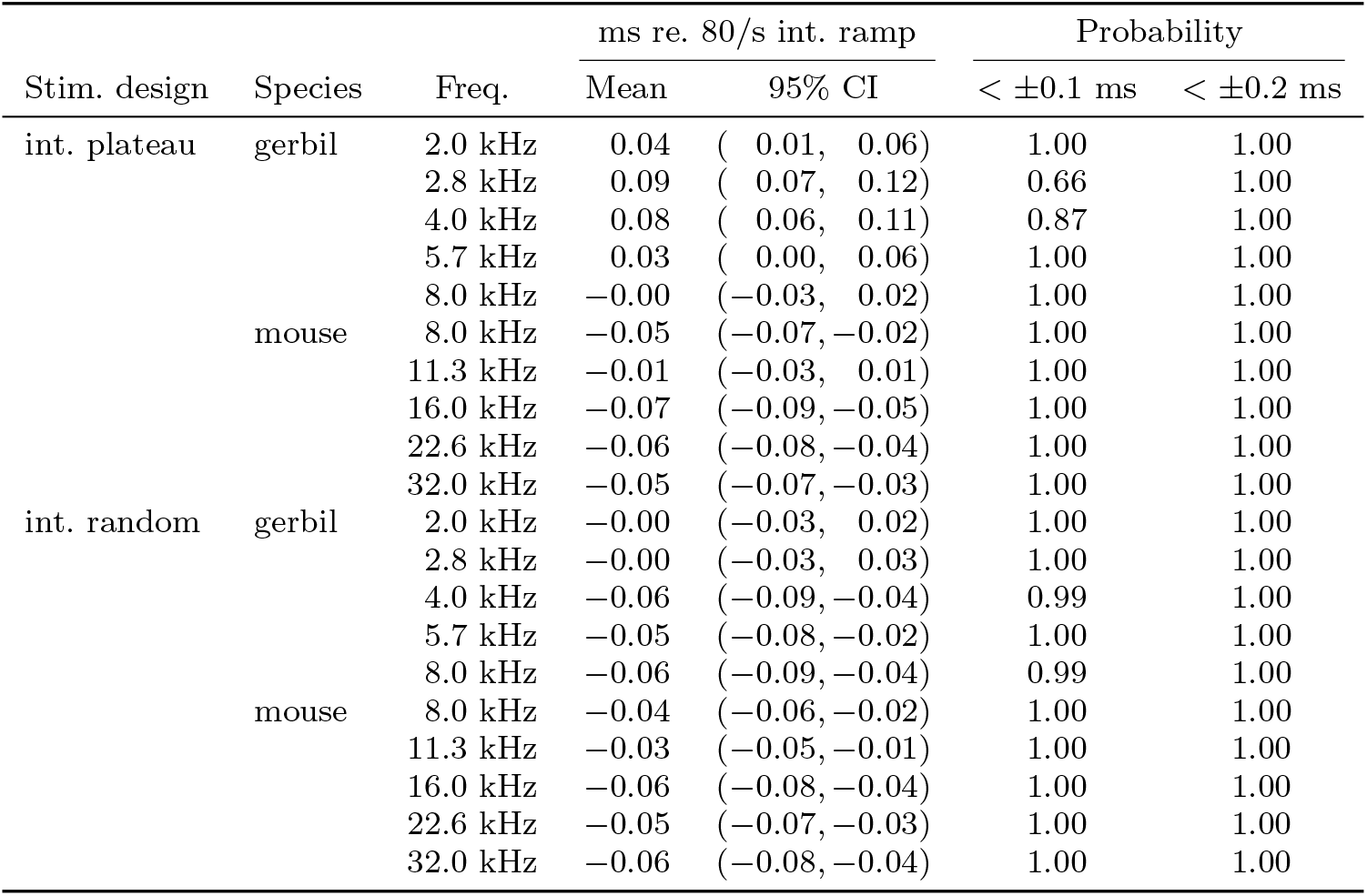
Change in wave 1 latencies for 80/s interleaved plateau and interleaved random stimulus designs in ms relative to 80/s interleaved ramp. Negative values indicate the latency was shorter than for 80/s interleaved ramp. Significance is assessed by the posterior probability that the difference in wave 1 latency from 80/s interleaved ramp is less than ±0.1 or ±0.2 ms.

Overall, we observe relatively small differences between the different interleaved configurations. Wave 1 amplitude is slightly larger for random compared to plateau and ramp, and threshold is slightly higher for random compared to plateau and ramp. Thus, among the three options, there is a trade-off between optimizing for threshold and wave amplitude.

### 3.4 ABR measurements in larger species

Experiments in larger species are often time-constrained to minimize the physiological and behavioral effects of prolonged anesthesia (Martin et al., 2014; Gottlieb et al., 2013; Sun et al., 2014). Thus, efficient ABR protocols may be especially beneficial for work with these animals. To test feasibility of the interleaved design in other species, we collected ABR data from ferret (*n* = 2 ears, Fig. 9A) and rhesus macaque (*n* = 4 ears, Fig. 9B). Average waveforms were clearly defined and permitted straightforward threshold measurement in both species. For macaque, data collected using a 60/s interleaved ramp design were compared to data collected using a 30/s conventional design (Fig. 9B). Due to time constraints in the macaque, we assessed a limited number of stimulus levels using the conventional design. In the ranges tested with both designs in macaque, results were comparable (Fig. 9C-E). Wave 1 amplitudes and latencies showed some variability across tone frequency (Fig. 9D-E). This variability likely results from the low number of ears and, because of limited data, the comparison of responses at 20 dB sensation level (SL), rather than more standard 80 dB SPL. Aside from these relatively small differences, the more rapid interleaved protocol produced results consistent with the slower conventional design.

**Fig. 9.**
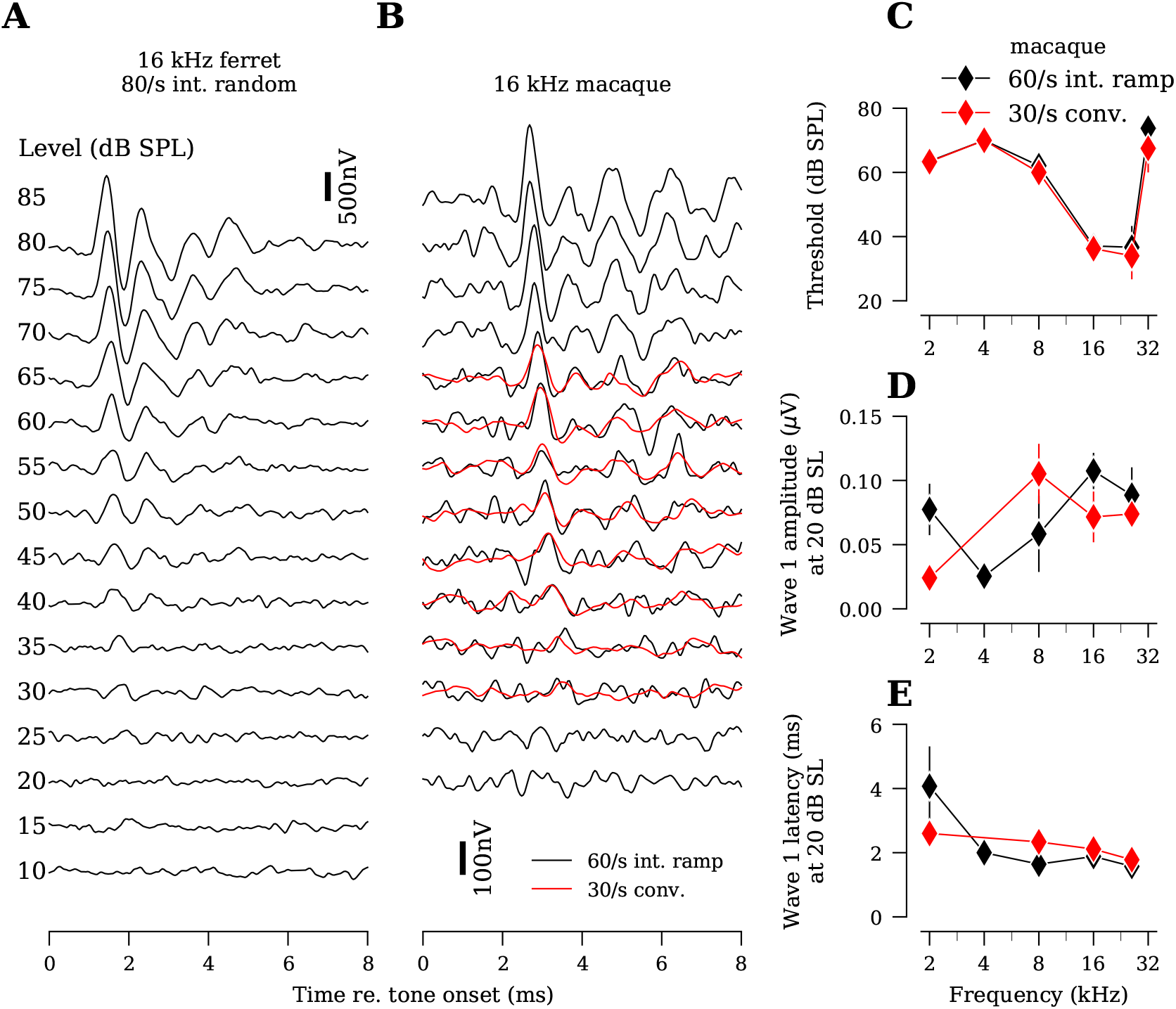
ABR data acquired using the interleaved stimulus design in other species. **A)** ABR waveforms from a single ferret ear using a 10 frequency (2 to 45.2 kHz), 15 level (10 to 80 dB SPL) 80/s interleaved random stimulus design. Data from 16 kHz are shown. **B)** ABR waveforms from a single rhesus macaque run on a 7 frequency (0.5 to 32 kHz), 14 level (20 to 85 dB SPL) 60/s interleaved ramp stimulus design. Overlaid are waveforms acquired using a single-frequency 30/s conventional approach. Data from 16 kHz are shown for both ferret and rhesus macaque. **C-E)** Average ABR thresholds, wave 1 amplitudes and wave 1 latencies from rhesus macaque (*n* = 4 ears), comparing 60/s interleaved ramp with 30/s conventional. Wave 1 amplitudes and latencies are assessed at 20 dB re ABR threshold (i.e., sensation level, dB SL). Error bars in all panels indicate ±SEM.

## 4 Discussion

We demonstrate that interleaving stimulus frequencies provides substantially more efficient auditory brainstem response (ABR) measurements than conventional designs. Interleaved stimuli reduce adaptation effects that can affect response amplitude measurements. The benefits are greatest when ABRs are required for multiple frequencies.

### Advantages of interleaved over conventional stimulus configurations

Interleaving tones of different frequencies reduces the effective rate at which individual frequencies are presented, thereby reducing adaptation to repeating tones. Thus, interleaved tones can be presented at faster rates without the same amount of adaptation as would be encountered with a conventional design. Specifically, we found:

- Interleaving five frequencies at 40 tones/s results in wave 1 amplitudes 25% larger than a conventional approach at 40 tones/s with no difference in acquisition time.
- Interleaving five frequencies at 80 tones/s produces wave 1 amplitudes comparable to a conventional approach at 40 tones/s while cutting acquisition time by 50% (i.e., when interleaving, two times as many frequencies can be acquired in the same amount of time as the conventional design).
- Interleaving five frequencies at 50 tones/s produces wave 1 amplitudes equivalent to a conventional approach at 10 tones/s while cutting acquisition time by 80% (i.e., when interleaving, five times as many frequencies can be acquired in the same amount of time as the conventional design).
- The interleaved stimulus design allows us to assess a larger range of frequencies and levels in time-sensitive experiments in rhesus macaque and ferret.

Differences in wave latency were clinically insignificant (e.g., less than 0.2 ms, Hwang et al., 2008; Musiek et al., 1989; Bauch et al., March-April 1982).

While our initial focus was on comparing the interleaved ramp to the conventional design, we found that the interleaved random design produces wave 1 amplitudes up to 38% larger than the interleaved ramp (Fig. 7D, Table 8). We did not directly compare interleaved random with conventional, but comparing Fig. 4D with Fig. 7D suggests we should see larger wave 1 amplitudes using a 80/s interleaved random stimulus design relative to 40/s conventional. However, the increased response amplitude for the interleaved random design comes with the trade-off of a small threshold elevation (1.9-4.3 dB) compared to the interleaved ramp design.

Later waves in the ABR are used to assess central auditory processing (Melcher and Kiang, 1996; Melcher et al., 1996). Amplitudes for the later waves were significantly enhanced by the interleaved stimulus design as compared to conventional when matched for rate (Fig. 5). This suggests that reduced adaptation of auditory nerve responses leads to greater activation of central auditory nuclei. Thus, experiments assessing later waves may also benefit from the interleaved stimulus design.

### Parallels with other studies

A study comparing a 9 tones/s conventional stimulus design with an 83 tones/s four-frequency interleaved ramp stimulus design (1 octave frequency spacing, 10 dB level spacing) found no significant difference in ABR thresholds, wave amplitudes or latencies (Mitchell et al., 1996). A follow-up study with a 100 tones/s seven-frequency interleaved ramp stimulus design (5 dB level spacing) found no difference in ABR thresholds, but showed slightly reduced wave 1 amplitudes in the interleaved ramp compared to conventional (Mitchell et al., 1999). Although their data suggest that one can go up to 83 tones/s without adaptation, they introduced an inter-train interval, reducing the average presentation rate from 83 to 59 tones/s (108 ms interval) and 100 to 88 tones/s (88 ms interval), respectively. In our study, there was no inter-train interval, thus these studies are consistent with our data (Fig. 6). Taken together, 50 to 59 tones/s likely represents the upper limit at which one can acquire data using an interleaved stimulus design while minimizing adaptation.

Traditionally, the maximum presentation rate for any ABR design is set by the analysis window used to measure evoked responses. For example, if the analysis window is 10 ms, the maximum presentation rate is 100 tones/s (10 ms separation between tone onsets). Faster tone presentations result in multiple tones falling in the analysis window. Recently, a study tested an approach in which tone pips of five frequencies are presented in parallel to each ear (Polonenko and Maddox, 2019). The interval between tone pips was randomized using a Poisson process with an average presentation rate of 200 tones/s across all frequencies. Since this study was performed in humans, only wave 5 was analyzed. These high presentation rates resulted in a reduction of up to 50% in wave 5 amplitudes and an increase in wave 5 latencies compared to an approach where a single frequency was presented at an average rate of 40 tones/s. Thus there may be faster approaches to acquiring ABRs, but they come with the trade-off of reduced wave amplitudes and increases in latencies.

The 5 ms tone burst with 0.5 ms rise-fall ramp used in this study is widely used in rodent studies (e.g., Kujawa and Liberman, 2006; Buran, 2009). In contrast, stimuli for human ABRs are of shorter duration with a slower rise-fall ramp (e.g., Polonenko and Maddox, 2019; Bramhall et al., 2017; Hood, 1998, Fig. 10). Although we did not assess how stimuli used in human ABRs might alter our conclusions, similar work in humans suggests that there is still a benefit of the interleaved stimulus design (Henry et al., 2000).

**Fig. 10.**
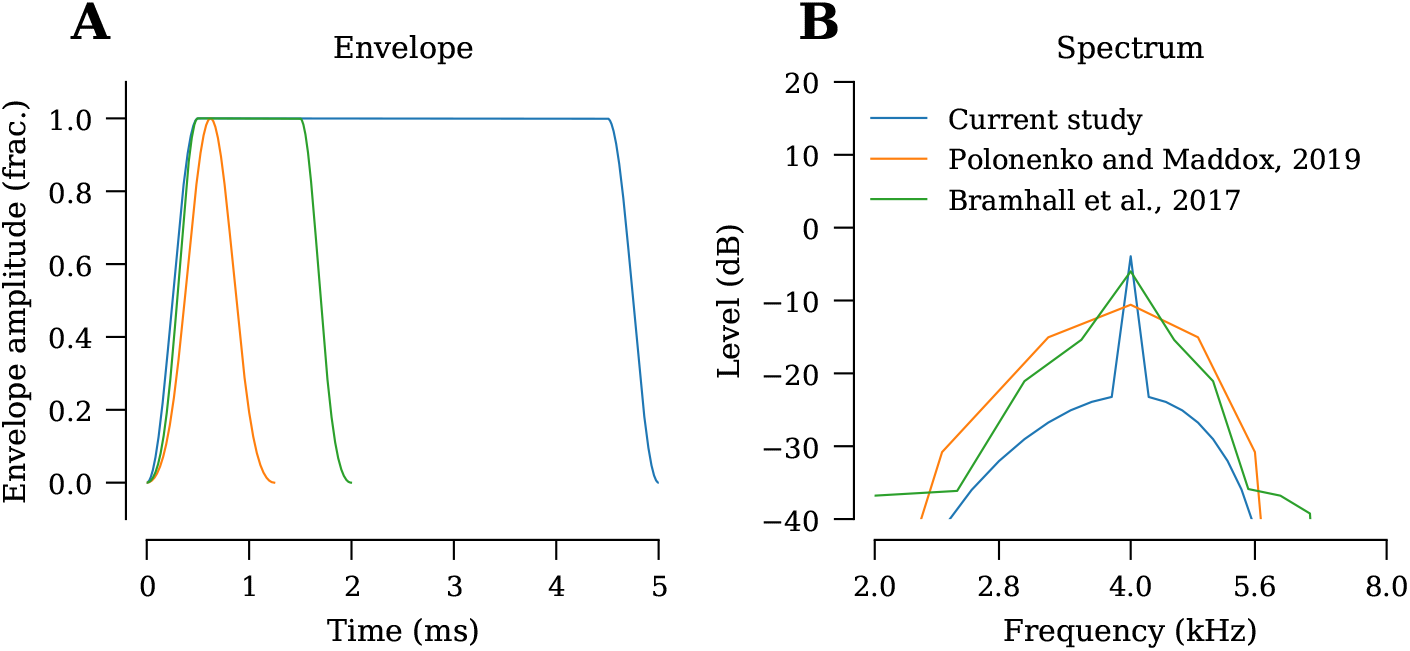
Comparison of the 4 kHz tone-burst stimulus used in this study with 4 kHz tone-burst stimuli used in human studies showing the envelope (**A**) and spectrum (**B**) of each stimulus.

### Interleaved configurations and mechanisms of adaptation

The phenomenon of forward masking limits the rate at which the ABR can be measured without affecting response amplitude (Spoor and Eggermont, 1971; Harris and Dallos, 1979; Burkard and Voigt, 1990). Here, the response to a tone pip can be partially masked by preceding tone pips. The different interleaved configurations determine which specific tone frequencies and levels form the masker for each response.

Based on the forward masking recovery equation developed by Harris and Dallos (1979), repetition rates of 10, 40, 50 and 80 tones/s suppress auditory nerve fiber activity by 0, 6, 9 and 15%, respectively. However, the actual suppression will be greater due to the cumulative effect of multiple tone pips acting as the masker. The reduced adaptation in the interleaved stimulus designs, compared to conventional at the same presentation rate, is likely due to the increased recovery time between tones with the same frequency and level. For example, at 40 tones/s using 5 frequencies and 12 levels, an 80 dB SPL, 4 kHz tone is presented only once every 1.5 s.

When considering the spacing of same-frequency masking tones, there is a caveat to consider with the interleaved ramp design. In this design, tones are grouped by frequency (Fig. 3B). Tones at the same frequency are presented in rapid succession from low to high intensity. Considering an 80 dB SPL tone is immediately preceded by a 75 dB SPL tone in the interleaved ramp, it might be surprising that we saw an increase in wave 1 amplitudes relative to the conventional stimulus design at the same presentation rate. However, lower level maskers produce less suppression than high level maskers (Spoor and Eggermont, 1971; Harris and Dallos, 1979), thereby resulting in a relatively small cumulative masking effect. The interleaved plateau design groups tones by level (Fig. 3C), increasing the interval between tones of the same frequency. The interval is proportional in size to the number of frequencies tested; e.g., in a five-frequency 80 tones/s interleaved plateau design, each frequency appears at a rate of 16 tones/s for an interstimulus interval of 62.5 ms. Despite this modified ordering, there was no difference in wave 1 amplitude between the ramp and plateau designs, potentially due to the spread of excitation at high stimulus levels resulting in a moderate level of masking comparable to that seen in the interleaved ramp design (Delgutte, 1996).

The interleaved random design had wave 1 amplitudes that were larger than for the interleaved ramp; however, there was a small increase in ABR threshold of approximately 1.9-4.3 dB (Table 7). This is likely due to high-intensity tones occasionally preceding low-intensity tones in the random sequence, in which case forward masking may suppress the neural response to the low-intensity tone and increase threshold. The lack of a commensurate reduction in wave 1 amplitude, despite the increase in threshold, is likely due to forward masking having a greater effect on low intensities than high intensities (Spoor and Eggermont, 1971).

All the interleaved stimulus designs offer an additional advantage over the conventional stimulus design. An animal’s anesthetic plane can fluctuate during an experiment, and this fluctuation may adversely affect a subset of frequencies and levels in the conventional stimulus design. In contrast, interleaved stimulus designs average out these fluctuations since all frequencies and levels are interrogated throughout the duration of the experiment.

### Limitations and alternatives

Shortcuts are sometimes taken when acquiring data using a conventional design. For example, the stimulus can be advanced in 5 dB increments from low to high level. Once threshold is identified, the experiment can either be halted or, if ABR amplitude is also desired, immediately advanced to the highest stimulus level. If shortcuts are routinely taken during conventional acquisition, the time-savings of switching to an interleaved design may not be as dramatic as reported here.

Similar shortcuts may be available for the interleaved stimulus design, but the efficacy of such approaches would need to be validated. The small differences in threshold, response amplitude and response latency between the three interleaved stimulus designs (Fig. 7) indicate that the ABR is sensitive to details of the stimulus train. Thus, time-saving shortcuts that alter the stimulus train (e.g., by dropping levels once certain criteria are met) may alter the response parameters of interest.

Although we used a fixed number of averages as the stopping criterion in our study, ABRs in humans are sometimes acquired until a particular residual noise or signal-to-noise (SNR) criterion is reached, particularly in clinical assessments (Norrix and Velenovsky, 2018; The Joint Committee on Infant Hearing, 2019). Audiologists typically perform a threshold search in which they rapidly iterate through different stimulus levels and frequencies to characterize the patient’s hearing loss. Under such scenarios, they are not interested in wave amplitude (although greater wave amplitude can enhance SNR). Since the interleaved stimulus design is constructed to maximize wave amplitude while minimizing changes in threshold, other approaches which focus on maximizing SNR may be more appropriate for assessment of auditory thresholds (e.g., Polonenko and Maddox, 2019; Valderrama et al., 2012; Burkard, 1991).

Since presenting ABR stimuli at high rates can be used to detect some neuropathologies (Hood, 1998; Burkard et al., 2007), any modification of the conventional approach, regardless of whether it’s a different averaging method, stimulus order or inter-trial interval, must be validated in both normal and hearing-impaired subjects (The Joint Committee on Infant Hearing, 2019).

We have demonstrated that the interleaved stimulus design offers a number of advantages over conventional approaches. However, the optimal set of parameters for the interleaved stimulus design (e.g., presentation rate) will likely depend on the animal species and hearing loss model as well as the stimulus levels and stimulus frequencies used. Since the interleaved stimulus design is not widely used, it is important to validate the results acquired with the interleaved stimulus design with the conventional approach when testing new species and forms of hearing loss.

## 5 Supplement

### 5.1 Hardware

Acoustic stimuli are typically generated digitally and converted to an analog waveform using digital to analog converters (DACs) in the data acquisition system (e.g., internal sound card or National Instruments card). Thus, the resolution of the DAC is important for handling stimulus waveforms that contain both low and high intensity tones. As described by Kester (2005), the theoretical dynamic range (i.e., signal-to-noise) ratio for converters is 6.02*N* + 1.76*dB* where *N* is the number of bits. This equation is commonly simplified to 6*N*. Regardless of which form of the equation is used, the calculation represents dynamic range under ideal conditions. The effective dynamic range is likely lower due to additional losses (e.g., wiring and interference from nearby electrical devices).

The 12-bit DAC used by Mitchell et al. (1996) has a theoretical maximum dynamic range of 74 dB. Although a standard ABR experiment may require only 70 dB of dynamic range for a given frequency, the calibration required by some closed-field speakers used in animal studies can vary by up to 20 dB across the frequency range of interest due to resonances shaped by the acoustic cavity (Hancock et al., 2015), requiring DACs that support dynamic ranges of at least 90 dB. Further, the effective dynamic range of the DAC is lower than the theoretical dynamic range. For example, modern 24-bit DACs have a theoretical dynamic range of 146 dB, but ambient noise limits it to approximately 120 dB in practice (Fujimori et al., 2000). Fortunately, 120 dB of dynamic range is sufficient for implementing the interleaved stimulus design across a large range of levels and frequencies. Therefore, equipment with 24-bit DACs is highly recommended when implementing the interleaved stimulus design.

### 5.2 Software

The software used in this study, psiexperiment, is available under the BSD three-clause license on Github and is free for anyone to download and modify. It currently is designed to work with National Instruments hardware and has been tested on the PXI system configuration recommended by Eaton-Peabody Laboratories (Hancock et al., 2015). Instructions for installing the software and configuring it to run the various stimulus protocols are posted on the website. Although other hardware platforms are not supported as of publication time, the modular nature of psiexperiment will simplify the process of getting it to run on other hardware platforms (e.g., high-quality 24-bit sound cards, TDT System 3, etc.).

## 6 Tables

## Acknowledgements

The authors would like to acknowledge Jesyin Lai’s assistance in acquiring data from ferret and Garnett McMillian for tutorials on implementing Bayesian models. This work was supported by the following grants:

– NIDCD R21 DC016969 to BNB

– NIDCD R01 DC014160 to JVB

– NIDCD R01 DC014950 to SVD

– Hearing Health Foundation Emerging Research Grant to BNB

– Pilot Program support from the Oregon National Primate Research Center, NIH award P51 OD011092.

## References

Bauch C, Rose D, Harner S (March-April 1982) Auditory Brain Stem Response Results from 255 Patients with Suspected Retrocochlear Involvement. Ear and Hearing 3(2):83–86

Bramhall NF, Konrad-Martin D, McMillan GP, Griest SE (2017) Auditory Brainstem Response Altered in Humans With Noise Exposure Despite Normal Outer Hair Cell Function. Ear and hearing 38(1):e1–e12, DOI 10.1097/AUD.0000000000000370

Buran BN (2009) Precision and reliability of cochlear nerve response in mice lacking functional synaptic ribbons. PhD thesis, Massachusetts Institute of Technology

Buran BN (2015) Auditory-wave-analysis: V1.1 — Zenodo. https://zenodo.org/record/17365

Buran BN, David SV (2018) ψ - psiexperiment. Zenodo, DOI 10.5281/zenodo.1405144

Burkard R (1991) Human brain-stem auditory evoked responses obtained by cross correlation to trains of clicks, noise bursts, and tone bursts. The Journal of the Acoustical Society of America 90(3):1398–1404, DOI 10.1121/1.401931

Burkard R, Voigt HF (1990) Stimulus dependencies of the gerbil brain-stem auditory-evoked response (BAER). III: Additivity of click level and rate with noise level. The Journal of the Acoustical Society of America 88(5):2222–2234, DOI 10.1121/1.400119

Burkard R, Shi Y, Hecox KE (1990) A comparison of maximum length and Legendre sequences for the derivation of brain-stem auditory-evoked responses at rapid rates of stimulation. The Journal of the Acoustical Society of America 87(4):1656–1664, DOI 10.1121/1.399413

Burkard RF, Eggermont JJ, Don M (2007) Auditory Evoked Potentials: Basic Principles and Clinical Application. Lippincott Williams & Wilkins

Cederholm JME, Froud KE, Wong ACY, Ko M, Ryan AF, Housley GD (2012) Differential actions of isoflurane and ketamine-based anaesthetics on cochlear function in the mouse. Hearing Research 292(1–2):71–79, DOI 10.1016/j.heares.2012.08.010

Delgutte B (1996) Psychophysiological Models. In: Fay RR, Popper AN (eds) Auditory Computation, Springer Handbook of Auditory Research, vol 6, Springer-Verlag, New York, pp 157–220

Dille MF, Ellingson RM, McMillan GP, Konrad-Martin D (2013) ABR Obtained from Time-Efficient Train Stimuli for Cisplatin Ototoxicity Monitoring. Journal of the American Academy of Audiology 24(9):769–781, DOI 10.3766/jaaa.24.9.2

Eysholdt U, Schreiner C (1982) Maximum length sequences – a fast method for measuring brain-stem-evoked responses. Audiology: Official Organ of the International Society of Audiology 21(3):242–250

Fausti SA, Mitchell CR, Frey RH, Henry JA, O’Connor JL (1994) Multiple-stimulus method for rapid collection of auditory brainstem responses using high-frequency (> or = 8 kHz) tone bursts. Journal of the American Academy of Audiology 5(2):119–126

Fowler CG, Noffsinger D (1983) Effects of stimulus repetition rate and frequency on the auditory brainstem response in normal cochlear-impaired, and VIII nerve/brainstem-impaired subjects. Journal of Speech and Hearing Research 26(4):560–567, DOI 10.1044/jshr.2604.560

Fujimori I, Nogi A, Sugimoto T (2000) A multibit delta-sigma audio DAC with 120-dB dynamic range. IEEE Journal of Solid-State Circuits 35(8):1066–1073, DOI 10.1109/4.859495

Furman AC, Kujawa SG, Liberman MC (2013) Noise-induced cochlear neuropathy is selective for fibers with low spontaneous rates. Journal of Neurophysiology 110(3):577–586, DOI 10.1152/jn.00164.2013

Gelman A, Carlin JB, Stern HS, Dunson DB, Vehtari A, Rubin DB (2013) Bayesian Data Analysis, 3rd edn. Chapman and Hall/CRC, Boca Raton

Gottlieb DH, Capitanio JP, McCowan B (2013) Risk factors for stereotypic behavior and self-biting in rhesus macaques (Macaca mulatta): Animal’s history, current environment, and personality. American Journal of Primatology 75(10):995–1008, DOI 10.1002/ajp.22161

Hancock K, Stefanov-Wagner IJ, Ravicz ME, Liberman MC (2015) The Eaton-Peabody Laboratories Cochlear Function Test Suite. In: Association for Research in Otolaryngology 38th Annual Midwinter Meeting, Baltimore, MD

Harris DM, Dallos P (1979) Forward masking of auditory nerve fiber responses. Journal of Neurophysiology 42(4):1083–1107

Henry JA, Fausti SA, Kempton JB, Trune DR, Mitchell CR (2000) Twenty-stimulus train for rapid acquisition of auditory brainstem responses in humans. Journal of the American Academy of Audiology 11(2):103–113

Hoffman MD, Gelman A (2014) The No-U-Turn Sampler: Adaptively Setting Path Lengths in Hamiltonian Monte Carlo. Journal of Machine Learning Research 15:1593–1623

Hood LJ (1998) Clinical Applications of the Auditory Brainstem Response. Singular Publishing Group

Hwang JH, Chao JC, Ho HC, Hsiao SH (2008) Effects of Sex, Age and Hearing Asymmetry on the Interaural Differences of Auditory Brainstem Responses. Audiology and Neurotology 13(1):29–33, DOI 10.1159/000107468

Kester W (ed) (2005) The Data Conversion Handbook. Newnes, Englewood Cliffs, NJ

Kiang NYS (1965) Discharge Patterns of Single Fibers in the Cat’s Auditory Nerve. MIT Press, Cambridge, MA

Kruschke JK (2013) Bayesian estimation supersedes the t test. Journal of Experimental Psychology General 142(2):573–603, DOI 10.1037/a0029146

Kujawa SG, Liberman MC (2006) Acceleration of Age-Related Hearing Loss by Early Noise Exposure: Evidence of a Misspent Youth. J Neurosci 26(7):2115–2123, DOI 10.1523/JNEUROSCI.4985-05.2006

Kujawa SG, Liberman MC (2009) Adding Insult to Injury: Cochlear Nerve Degeneration after “Temporary” Noise-Induced Hearing Loss. The Journal of neuroscience : the official journal of the Society for Neuroscience 29(45):14077–14085, DOI 10.1523/JNEUROSCI.2845-09.2009

Lentz JJ, Gordon WC, Farris HE, MacDonald GH, Cunningham DE, Robbins CA, Tempel BL, Bazan NG, Rubel EW, Oesterle EC, Keats BJ (2010) Deafness and Retinal Degeneration in A Novel USH1C Knock-In Mouse Model. Developmental neurobiology 70(4):253–267, DOI 10.1002/dneu.20771

Lin HW, Furman AC, Kujawa SG, Liberman MC (2011) Primary Neural Degeneration in the Guinea Pig Cochlea After Reversible Noise-Induced Threshold Shift. JARO: Journal of the Association for Research in Otolaryngology 12(5):605–616, DOI 10.1007/s10162-011-0277-0

Martin LD, Dissen GA, McPike MJ, Brambrink AM (2014) Effects of Anesthesia with Isoflurane, Ketamine, or Propofol on Physiologic Parameters in Neonatal Rhesus Macaques (Macaca mulatta). Journal of the American Association for Laboratory Animal Science : JAALAS 53(3):290–300

McMillan GP, Cannon JB (2019) Bayesian Applications in Auditory Research. Journal of speech, language, and hearing research: JSLHR 62(3):577–586, DOI 10.1044/2018_JSLHR-H-ASTM-18-0228

Melcher JR, Kiang NY (1996) Generators of the brainstem auditory evoked potential in cat III: Identified cell populations. Hearing Research 93(1-2):52–71, DOI 10.1016/0378-5955(95)00200-6

Melcher JR, Knudson IM, Fullerton BC, Guinan Jr JJ, Norris BE, Kiang NY (1996) Generators of the brainstem auditory evoked potential in cat. I. An experimental approach to their identification. Hearing Research 93(1–2):1–27, DOI 10.1016/0378-5955(95)00178-6

Millan J, Ozdamar O, Bohórquez J (2006) Acquisition and analysis of high rate deconvolved auditory evoked potentials during sleep. Conference proceedings: Annual International Conference of the IEEE Engineering in Medicine and Biology Society IEEE Engineering in Medicine and Biology Society Annual Conference 1:4987–4990, DOI 10.1109/IEMBS.2006.260399

Mitchell C, Kempton JB, Creedon T, Trune D (1996) Rapid acquisition of auditory brainstem responses with multiple frequency and intensity tonebursts. Hearing Research 99(1):38–46, DOI 10.1016/S0378-5955(96)00081-0

Mitchell CR, Kempton JB, Creedon TA, Trune DR (1999) The Use of a 56-Stimulus Train for the Rapid Acquisition of Auditory Brainstem Responses. Audiology and Neurotology 4(2):80–87, DOI 10.1159/000013824

Mitchell CR, Ellingson RM, Henry JA, Fausti SA (2004) Use of auditory brainstem responses for the early detection of ototoxicity from aminoglycosides or chemotherapeutic drugs. Journal of Rehabilitation Research and Development 41(3A):373–382, DOI 10.1682/jrrd.2003.05.0089

Mouney DF, Cullen JK, Gondra MI, Berlin CI (1976) Tone burst electrocochleography in humans. Transactions Section on Otolaryngology American Academy of Ophthalmology and Otolaryngology 82(3 Pt 1):348–355

Musiek F, Johnson G, Gollegly K, Josey A, Glasscock M (1989) The Auditory Brain Stem Response Interaural Latency Difference (ILD) in Patients with Brain Stem Lesions. Ear and Hearing 10(2):131–134

Norrix LW, Velenovsky D (2018) Clinicians’ Guide to Obtaining a Valid Auditory Brainstem Response to Determine Hearing Status: Signal, Noise, and Cross-Checks. American Journal of Audiology 27(1):25–36, DOI 10.1044/2017_AJA-17-0074

Nuzzo R (2014) Scientific method: Statistical errors. Nature News 506(7487):150, DOI 10.1038/506150a

Paludetti G, Maurizi M, Ottaviani F (1983) Effects of stimulus repetition rate on the auditory brain stem responses (ABR). The American Journal of Otology 4(3):226–234

Polonenko MJ, Maddox RK (2019) The Parallel Auditory Brainstem Response. Trends in Hearing 23:2331216519871395, DOI 10.1177/2331216519871395

Robles L, Ruggero MA (2001) Mechanics of the Mammalian Cochlea. Physiological reviews 81(3):1305–1352

Ruebhausen MR, Brozoski TJ, Bauer CA (2012) A comparison of the effects of isoflurane and ketamine anesthesia on auditory brainstem response (ABR) thresholds in rats. Hearing Research 287(1):25–29, DOI 10.1016/j.heares.2012.04.005

Salvatier J, Wiecki TV, Fonnesbeck C (2016) Probabilistic programming in Python using PyMC3. PeerJ Computer Science 2:e55, DOI 10.7717/peerj-cs.55

Smith DI, Mills JH (1989) Anesthesia effects: Auditory brain-stem response. Electroencephalography and Clinical Neurophysiology 72(5):422–428, DOI 10.1016/0013-4694(89)90047-3

Spoor A, Eggermont JJ (1971) Action potentials in the cochlea. Masking, adaptation and recruitment. Audiology: Official Organ of the International Society of Audiology 10(5):340–352

Stamper GC, Johnson TA (2015) Auditory function in normal-hearing, noise-exposed human ears. Ear and hearing 36(2):172–184, DOI 10.1097/AUD.0000000000000107

Sun L, Li Q, Li Q, Zhang Y, Liu D, Jiang H, Pan F, Yew DT (2014) Chronic ketamine exposure induces permanent impairment of brain functions in adolescent cynomolgus monkeys. Addiction Biology 19(2):185–194, DOI 10.1111/adb.12004

Szucs D, Ioannidis JPA (2017) When Null Hypothesis Significance Testing Is Unsuitable for Research: A Reassessment. Frontiers in Human Neuroscience 11, DOI 10.3389/fnhum.2017.00390

The Joint Committee on Infant Hearing (2019) Year 2019 Position Statement: Principles and Guidelines for Early Hearing Detection and Intervention Programs. The Journal of Early Hearing Detection and Intervention 4(2):1–44

Valderrama JT, Alvarez I, de la Torre A, Carlos Segura J, Sainz M, Luis Vargas J (2012) Recording of auditory brainstem response at high stimulation rates using randomized stimulation and averaging. The Journal of the Acoustical Society of America 132(6):3856–3865, DOI 10.1121/1.4764511

